# Fractionating difficulty during sentence comprehension using functional neuroimaging

**DOI:** 10.1101/2023.08.09.552675

**Authors:** Malathi Thothathiri, Jana Basnakova, Ashley G. Lewis, Josephine M. Briand

**Author notes:** Address all correspondence to: Malathi Thothathiri Department of Speech, Language and Hearing Sciences The George Washington University Washington, DC 20052.

## Abstract

Sentence comprehension is highly practiced and largely automatic, but this belies the complexity of the underlying processes. We used functional neuroimaging (fMRI) to investigate garden-path sentences that cause difficulty during comprehension, in order to unpack the different processes used to support sentence interpretation. By investigating garden-path and other types of sentences within the same individuals, we functionally profiled different regions within the temporal and frontal cortices in the left hemisphere. The results revealed that different aspects of comprehension difficulty are handled by left posterior temporal, left anterior temporal, ventral left frontal, and dorsal left frontal cortices. The functional profiles of these regions likely lie along a spectrum of specificity to generality, including language-specific processing of linguistic representations, more general conflict resolution processes operating over linguistic representations, and processes for handling difficulty in general. These findings suggest that difficulty is not unitary and that there is a role for a variety of linguistic and non-linguistic processes in supporting comprehension.

## Introduction

The ability to construct and understand sentences (*cf.* individual sounds or symbols) is an essential feature that separates human language from other forms of human and non-human communication. Most sentences are not stored and reproduced in whole. Every day, we generate new sentences to communicate ideas and thoughts, and understand new sentences produced by others. Although highly practiced, sentence comprehension is a complex ability that is supported by multiple processes, such as retrieving words from the mental lexicon, building syntactic structures, and resolving conflict between competing representations. The role of these different processes can be determined by studying sentences that are more difficult to process than the simplest cases. So-called “garden-path” sentences have a long history in psycholinguistics and neurolinguistics because they allow researchers to separate different linguistic processes (syntax, semantics, pragmatics) and also to determine how broader functions like cognitive control and working memory assist comprehension (Christianson et al., 2001; den Ouden et al., 2016; Just & Carpenter, 1992; Novick et al., 2005). Consider a garden-path sentence like (1):

### (1) As the men wrestled the rivals arrived at the gymnasium

As readers and listeners process the sentence, they initially tend to interpret “rivals” as the object of wrestling (i.e., that the men wrestled the rivals). However, upon encountering “arrived” and the subsequent words, the language comprehension system experiences conflict with this interpretation, which should trigger syntactic re-analysis and re-interpretation (i.e., that the men were wrestling someone else and not the rivals, who just arrived). This kind of tripping-up makes garden-path sentences more difficult than say, a sentence like “The cat sat on the mat.” But what exactly is “difficulty”? Is it a unitary concept, or are different aspects of a difficult comprehension situation handled by different neural systems? In this paper, we address this issue by measuring neural activation for different types of sentences, which were designed to isolate the contribution of different processes, in the same set of individuals.

Prior literature has identified a key set of processes that could be recruited to different extents for different types of sentences. These include syntactic processing, semantic/pragmatic processing, cognitive control, and working memory. Historically, syntax was considered the predominant process that is used for sentence comprehension because the meaning of a sentence ultimately depends on its syntactic structure. Syntax allows us to differentiate between “The dog chased the cat” and “The cat chased the dog” despite the two sentences containing the same words. However, several studies have now shown that semantics and pragmatics can also exert a powerful influence on sentence interpretation, sometimes even overriding the meaning indicated by syntax (see e.g., Altmann & Kamide, 1999; Ferreira, 2003; Kim & Osterhout, 2005). For example, a sentence like “The dog was bitten by the man” may be interpreted instead as the dog biting the man due to our semantic and pragmatic knowledge about the world. Cognitive control may be relevant for supporting accurate comprehension especially in demanding situations such as ambiguous sentences, conflicting interpretations, and potentially noisy input conditions (Novick et al., 2005; Thothathiri et al., 2012a; Peelle & Wingfield, 2022). Working memory is relevant for integrating incoming words into a cohesive structural interpretation (Just & Carpenter, 1992; Lewis et al., 2006).

In functional neuroimaging (fMRI) studies, contrasts between different conditions that vary on a particular dimension are used to identify the neural substrates associated with a particular process. For example, syntactically complex sentences may be compared to syntactically simpler sentences in an attempt to locate regions that are relevant to syntactic processing. However, herein lies a challenge. Sentences that are difficult along one dimension are also usually difficult along other dimensions. For example, compare the garden-path sentence (1) above to a non-garden-path sentence like (2):

### (2) While the zookeeper fed the ponies the stallion stomped its hoof

For (1), syntactic re-analysis is needed in order to arrive at the correct interpretation (i.e., that the men did not necessarily wrestle the rivals). In contrast, in (2), the structure is unambiguous—it is clear that the zookeeper fed the ponies and not the stallion, and no re-analysis is needed. Thus, a contrast between (1) and (2) might be expected to identify regions involved in syntactic analysis. However, this is not the only possible differential process in this comparison: (1) may also trigger additional semantic/pragmatic processing to reconcile the different conflicting interpretations, cognitive control to resolve conflict, and working memory to re-process the sentence. Separating these different dimensions of difficulty is important for understanding the various sub-components of sentence processing and identifying the neural substrates of each component.

We used a functional profiling approach to fractionate the processing of difficult-to-comprehend sentences. Specifically, we examined how regions of interest (ROIs) implicated in processing garden-path versus non-garden-path sentences are activated for *other* sentences that vary in their syntactic re-analysis, semantic/pragmatic processing, cognitive control, and working memory demands. Below, we briefly review prior literature on the processes used for sentence comprehension in general and the literature on garden-path sentences in particular before describing the design of the present study and its contributions.

#### Prior Literature on Sentence Comprehension Processes

Sentences are composed of multiple words connected together by structural or syntactic rules. Accordingly, early neurolinguistic studies focused on identifying the locus of syntactic operations in the brain (see e.g., Caplan et al. (2001) and Dapretto & Bookheimer (1999) for a summary). Typical experimental designs contrasted sentences with different syntactic complexity, sentences containing syntactic versus other kinds of errors, or tasks that emphasized syntactic versus other kinds of processing (Caplan et al., 2001; Dapretto & Bookheimer, 1999; Embick et al., 2000). Many studies pointed towards a role for the left frontal cortex, especially Brodmann areas 44 and/or 45, collectively known as Broca’s area (Dapretto & Bookheimer, 1999; Embick et al., 2000). However, some early as well as more recent studies have documented the involvement of left posterior temporal and inferior parietal regions (Caplan et al., 2001; Thothathiri et al., 2012b; Wartenburger et al., 2004; Yokoyama et al., 2007). Damage to the temporo-parietal and not the frontal cortex predicts syntactic comprehension deficits in aphasia (Thothathiri et al., 2012b. See also Fridrikkson et al., 2018). Based on a meta-analysis of more than 35 neuroimaging (fMRI/PET) studies of sentence comprehension, Walenski et al. (2019) found evidence for an association between syntactic processing and both left frontal and left posterior temporal regions. Overall, there is growing consensus for the posterior temporal lobe’s involvement in syntactic processing during comprehension. For the frontal lobe, there is debate about whether its role may be best described as being more specific to sentence production than comprehension or broader resources like cognitive control or working memory (Matchin et al., 2020; Thothathiri et al., 2012a, 2012b; Walensky et al., 2019. See also Diachek et al., 2020).

Sentence comprehension is ultimately about understanding meaning. The meanings of individual words must be combined to compute the compositional meaning of the sentence. In contrast to the syntactic processing that is tied to more posterior temporal regions, semantic composition has been linked to the left anterior temporal lobe (ATL). Left ATL has been linked to semantic memory based on word-level evidence from semantic dementia, compositional processing at the level of phrases and sentences, and to interactions between word-level and phrasal-level information (Lambon Ralph et al., 2010; Westerlund and Pylkkanen, 2014). Although damage to this region is not routinely associated with sentence comprehension deficits, its involvement in semantic processing could make it relevant for language comprehension under some conditions (Walensky et al., 2019).

Sentence comprehension difficulty is not only about challenges in syntactic or semantic processing but could also be about broader cognitive resources like cognitive control and working memory. Cognitive control, or the ability to resolve conflict between competing representations, has been argued to be relevant especially for more difficult-to-comprehend sentences. Garden-path sentences like (1) above create conflict between the original interpretation (e.g., that the men wrestled the rivals) and other information from the sentence (e.g., that the rivals just arrived) and cognitive control can be useful for resolving such conflict. Therefore, at least some difficult sentences could be difficult due to their cognitive control demands. Sentence processing also requires storing and building structured representations from words as the sentence unfolds incrementally i.e., working memory (Lewis et al., 2006; Shain et al., 2022). For example, understanding a sentence like “We sang a song that our daughter really likes” requires holding on to the word “song” until it can be linked to the verb “likes” at the end (*cf. “*Our daughter really likes a song that we sang”). Thus, syntactically difficult sentences may require more working memory resources than their simpler counterparts. Prior studies have provided broad support for the role of both cognitive control and working memory in sentence comprehension, based on neuroanatomical co-localization of those functions with sentence processing and causal links between damage to or upregulation of those abilities and better sentence comprehension (e.g., Horne et al., 2022; Hsu et al., 2017; Thothathiri et al., 2018; Vuong & Martin, 2015; Ye & Zhou, 2009). The left frontal cortex has been linked to both cognitive control and working memory, as part of networks working in tandem with temporal, parietal, and medial frontal regions (e.g., see Botvinick et al., 2001; Hsu et al., 2017; Shain et al., 2022).

Different neurolinguistic frameworks of language processing all accord a role for the processes discussed above in sentence comprehension. But they differ in the weighting allocated to different components. Hagoort (2005)’s Memory, Unification and Control (MUC) model proposes that language processing utilizes representations stored in memory in the left temporal cortex. Unification of different linguistic representations is hypothesized to be coordinated by the left inferior frontal cortex. Last but not least, control operations in dorsolateral and medial frontal cortices support goal-directed use of language in different contexts. The MUC model explicitly allows for interactive, concurrent processing of different sources of information (e.g., syntactic, semantic and pragmatic). It does not prioritize syntax. It also allows for a role for domain-general control operations in language use. By comparison, Friederici (2002)’s neuroanatomical model of sentence processing proposes a syntax-first view wherein syntactic processes precede semantic processes initially and the two interact only during later phases. Semantic representations are hypothesized to be stored in the temporal cortex with the frontal cortex supporting controlled or strategic use of those representations. For syntax, the temporal cortex and the most inferior parts of the frontal cortex are thought to be relevant for syntactic operations with other less inferior parts of the frontal cortex engaged for working memory.

More recently, Fedorenko and colleagues have used a single-subject localization approach to argue that most of language processing occurs within a bilateral temporo-frontal language-selective network. They distinguish some lateral frontal areas within this network from other nearby lateral frontal regions that they argue are part of a more domain-general Multiple Demand (MD) network, which is not specific to language processing (Fedorenko et al., 2012; Diachek et al., 2020; Shain et al., 2022). Although many language tasks activate both the language and the MD networks, these authors have suggested that the latter is primarily engaged when there is an explicit secondary task going beyond passive sentence comprehension (Diachek et al., 2020). Thus, left frontal regions in this framework are split between those engaged in language-specific operations within the language network and those involved in more domain-general processes within the MD network.

To summarize, converging evidence in the field suggests that both temporal and frontal regions within the left hemisphere are involved in sentence comprehension. Temporal regions are widely thought to store the linguistic representations—syntactic and semantic—that are relevant for processing sentences. The precise role of the left frontal cortex is more debated. There is consensus that the more dorsal portions are involved in domain-general processes that support both linguistic and non-linguistic tasks. However, the more ventral/inferior portions have been linked variously to unification of syntactic and non-syntactic representations, syntax-specific operations, working memory, and cognitive control (Hagoort, 2005; Hsu et al., 2017; Fiebach et al., 2005; Friederici, 2002; Thothathiri et al., 2012a).

#### Prior Literature on Garden-Path Sentences

For garden-path sentences like (1), prior electrophysiological (ERP) studies have demonstrated a P600 signal at the point of disambiguation. Qian et al. (2018) found a P600 effect for garden-path versus non-garden-path sentences in healthy adults. Sheppard et al. (2017) contrasted garden-path sentences with plausible versus implausible noun phrases (e.g., *While the band played the song/beer…*) and found an N400-P600 complex for the latter compared to the former in healthy adults and patients with anomic aphasia. In ERP studies, the P600 signal is seen for a range of sentence stimuli, including those with syntactic violations, thematic role assignment errors, or even more broadly, any violation of expectations during sentence comprehension. This has led some to argue that it reflects conflict monitoring (van de Meerendonk et al., 2010). Together, a P600 effect during the processing of sentences similar to (1) confirms that the comprehension system can utilize disambiguating syntactic or semantic information to detect conflict between interpretations and undertake structural reanalysis relatively quickly. But it does not clarify the various processes used to understand these sentences and their neural correlates.

Two previous fMRI studies have examined the neural correlates of comprehending garden-path sentences like (1). Hsu et al. (2017) compared activation for reading garden-path sentences (e.g., “While the thief hid the jewelry sparkled”) and their non-garden-path counterparts (e.g., “While the thief hid, the jewelry sparkled” with the critical comma). They found that the top 100 voxels activated for this contrast in the left inferior frontal cortex were also activated for three other cognitive control tasks (Stroop, n-back with lures and recent negatives) and vice versa. This co-localization across tasks indicated that the left inferior frontal cortex was a general-purpose hub for cognitive control. At the same time, functional connectivity analyses revealed distinct connected areas for different tasks, indicating that specialization might arise at the network level rather than due to specialization within the frontal cortex per se.

den Ouden and colleagues (2016) conducted whole-brain and ROI analyses of garden-path and non-garden-path sentences presented auditorily. They found evidence suggesting a key role for the posterior temporal cortex (*cf.* frontal or anterior temporal cortex) in processing syntactic ambiguity. Within the frontal cortex, more fine-grained ROI analyses detected differences between sub-regions such that Brodmann area 45 responded only to sentences where re-analysis occurred and more posterior-dorsal Brodmann area 44 responded to all sentences with ambiguity/conflict. Overall, the findings accorded with prior literature (see above) in indicating a central role of the posterior temporal cortex in syntactic processing. For the frontal cortex, the effects were subtler and limited to cases of ambiguity or reanalysis.

Neither of the previous studies was designed to fractionate difficulty during sentence comprehension. den Ouden et al. (2016) focused on the interaction between prosodic and plausibility cues during auditory sentence comprehension. Hsu et al. (2017) did conduct a secondary analysis of MD network regions to distinguish between cognitive control and broad difficulty that bears on the interpretation of our results (see Discussion). However, their emphasis was on domain-generality rather than on understanding the set of processes used to comprehend difficult garden-path sentences. The present study was designed to address this gap and answer questions about whether difficulty in the context of sentence comprehension is handled by a variety of regions that support handling different aspects of that difficulty.

#### The present study

We sought to fractionate the difficulty associated with hard-to-comprehend garden-path sentences by comparing activation for different kinds of difficult sentences within the same regions of interest (ROIs). We used a sentence reading plus comprehension question task with five different types of sentences, including (1) and (2), reproduced from above. The first three sentence types contained a subordinate clause followed by a main clause. They were used to contrast garden-path and non-garden-path sentences (1 vs 2; 3 vs 2) and also two different variants of garden-path sentences (1 vs 3) in order to understand how the brain handles ambiguity and conflict during sentence comprehension. The last two sentence types (4 and 5) manipulated difficulty along a different dimension, namely working memory. They were included to help separate the neural substrates of processes that handle ambiguity and conflict (e.g., cognitive control for resolving conflict) from those underlying working memory. Below, we first discuss sentence types (1)-(3) and then (4)-(5).

(1) Implausible: *As the men wrestled the rivals arrived at the gymnasium*.
(2) Control: *While the zookeeper fed the ponies the stallion stomped its hoof*.
(3) Plausible: *While the farmer steered the green tractor pulled the plough*.
(4) Long WM: *Ezekiel bragged about his circus art skills and recorded himself playing a Beethoven sonata on the piano*.
(5) Short WM: *Natalie and Margaret called themselves an overpriced cab and visited the natural history museum*.

Control sentences like (2) did not contain syntactic ambiguity or conflict. By contrast, in sentences like (1), there was syntactic ambiguity before “arrived” because “the rivals” could be either the end of the subordinate clause that started with “As the men” or the beginning of a new main clause that started with “the rivals”. This should generate conflict because the parser originally constructs an analysis consistent with the former but then encounters “arrived”, which is inconsistent with that analysis. The same kind of ambiguity and conflict also existed in sentences like (3). Thus, comparisons between garden-path sentences like (1) and (3) versus non-garden-path sentences like (2) can identify regions involved in cognitive control for resolving conflict arising from temporary syntactic ambiguity, and syntactic analysis for reanalyzing the sentence.

The key difference between (1) and (3) was that the ending of the sentence in (1)—that the rivals arrived at the gymnasium—made the original misinterpretation—that the men wrestled the rivals—unlikely. Therefore, these sentences were labeled “Implausible”. In contrast, for (3), the ending of the sentence—that the green tractor pulled the plough—left the original interpretation—that the farmer steered the green tractor—still semantically viable. Therefore, these sentences were labeled “Plausible”. A number of previous studies have employed Implausible and Plausible sentences and shown that comprehenders are more likely to retain the original misinterpretation and say Yes to questions like “Did the farmer steer the tractor?” in the Plausible condition than to questions like “Did the men wrestle the rivals?” in the Implausible condition (see e.g., Christianson et al., 2001, 2006). Using terminology employed in this literature, Plausible sentences like (3) are more subject to “lingering misinterpretation” than Implausible sentences like (1) (Christianson et al., 2001; Slattery et al., 2013). Put the other way, comprehenders are more successful in letting go of the original misinterpretation for Implausible than Plausible sentences, suggesting that the former recruits some additional process or processes that can help reconcile incompatible interpretations. Thus, differential activation for the Implausible versus Plausible condition can identify regions involved in the additional process(es).

As mentioned above, we also included two other types of sentences that were both long and could induce comprehension difficulty along a different dimension, namely working memory. These sentences (Long and Short WM) contained reflexive pronouns (herself/himself/themselves) that had to be bound to an antecedent for comprehension (***Ezekiel*** *bragged about … and recorded **himself***). The distance separating the antecedent from the reflexive pronoun was longer for the long than the short WM condition. Importantly, while these sentences were expected to induce difficulty related to working memory demands, they did not contain ambiguity or conflict and were therefore not expected to recruit cognitive control or syntactic analysis regions more than the Control condition.

Additionally, we tested sentence reading in a non-linguistic context that manipulated visual attentional demands. Participants saw a sentence below a visual stimulus and were asked to indicate whether the sentence accurately described the image (Figure 1).

(6) More/Less: *Less than half of the pigs are to the right of the lamp*.
(7) All/Some: *Some of the sunglasses shown here are to the left of the lamp*.

**Figure 1.**
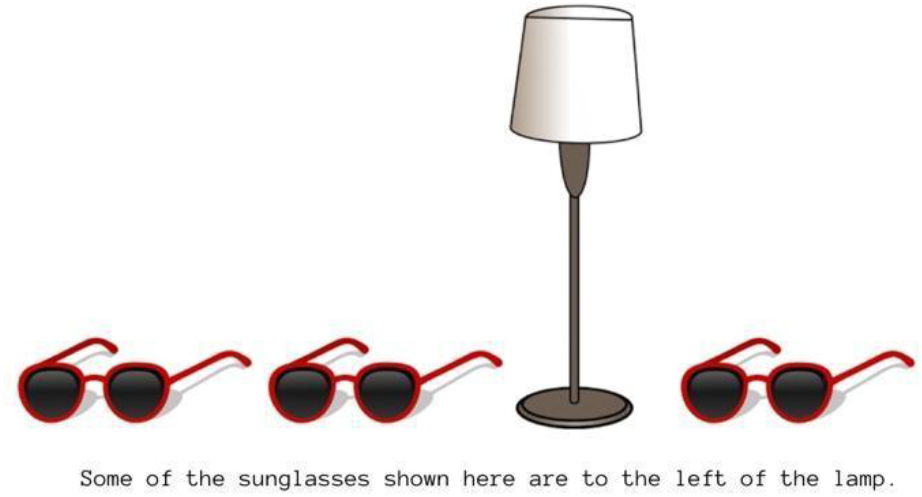
Example image for the Visual Attention task.

In the More/Less condition, the sentences contained the quantifiers “More than half” or “Less than half”. In the All/Some condition, the sentences contained the quantifiers “All” or “Some”. For More/Less, doing the task correctly required evaluating the total number of objects in the image and then determining whether more (or less) than half of that number was present on the indicated side (left/right). By contrast, for All/Some, the task could be completed more simply by assessing whether all of the objects were to one side of the lamp or whether they were distributed (in any way) on both sides. Thus, the More/Less condition was expected to be more difficult than All/Some but this difficulty is due to demands on visual attention rather than on linguistic processes like syntactic or semantic processing or broader processes associated with sentence processing such as cognitive control or working memory.

To summarize, we attempted to separate different sources of difficulty that can make comprehension challenging when sentences are not simple or straightforward. Garden-path sentences can trigger additional syntactic analysis, semantic/pragmatic processing, cognitive control, and potentially working memory and general task-demands relative to syntactically unambiguous sentences. Therefore, we tested the comprehension of sentences that tap these different resources to different extents, in the same set of individuals. We reasoned that Implausible and Plausible sentences would recruit additional cognitive control and syntactic analysis processes relative to Control because they contain ambiguity and generate conflict between interpretations. Further, Implausible sentences should trigger additional processes like deeper semantic/pragmatic processing compared to Plausible sentences in order to handle the implausibility of maintaining the original (mis)interpretation after encountering the information at the end of the sentence. If regions involved in working memory, separately from those engaged in cognitive control, are used to deal with garden-pathing, we should see increased recruitment for Long and Short WM in those specific regions. Finally, if there are regions that assist sentence comprehension via a broad response to task difficulty, we should see an effect of the visual attention manipulation (More/Less > All/Some) in those specific regions. In sum, by looking at the functional profile of regions engaged for implausible garden-path versus non-garden-path sentences, we sought to fractionate the neural correlates into different processes that can support the comprehension of difficult-to-understand sentences. See Table 1 for a summary of the logic.

**Table 1:** Logic of the Functional Profiling Approach.

| Implausible<br>> Control | Plausible ><br>Control | Long WM ><br>Control | Short WM ><br>Control | More/Less ><br>All/Some | ***Region is involved in*** |
| --- | --- | --- | --- | --- | --- |
| ✓ | ✓ | x | x | x | ***Cognitive control or syntactic reanalysis*** |
| ✓ | x | x | x | x | ***Semantic/pragmatic processing?*** |
| ✓ | ✓ | ✓ | ✓ | x | ***Working memory demands*** |
| ✓ | ✓ | x | x | ✓ | ***General task difficulty demands*** |

## Methods

### Norming Study

We normed the stimuli to be used in the study prior to collecting data. Thirty-seven adults (15 Male, 15 Female, 21-36 years, Median age=31^1^) participated in a web-based experiment. Participants were recruited using Prolific, a web-based platform for conducting research. Participants self-reported being right-handed, native English speakers. Prolific collects demographic information independently of specific studies, minimizing concerns about misrepresentation by the participants. Stimuli were presented using Python on Pavlovia (https://pavlovia.org).

Participants completed two tasks. In the sentence reading task, they read 42 sentences in each of the five conditions (Control, Implausible, Plausible, LongWM, ShortWM) word by word and answered yes/no comprehension questions. In the visual attention task, they saw a picture and read the sentence below it to answer whether the sentence accurately described the picture. There were 42 trials in the More/Less condition (21 More, 21 Less) and 42 trials in the All/Some condition (21 All, 21 Some). For both tasks, they used the “j” key for a yes and the “k” key for a no response.

For the sentence reading task, we confirmed that the Plausible sentences were more subject to lingering misinterpretation than the Implausible sentences (Plausible mean accuracy=59.5%. Implausible mean accuracy=79.6%. z=-5.13, p<.001). Accuracies for Long WM and Short WM sentences were at ceiling and did not differ from one another (97% for both. z=.56, p=.57). For the Control condition, accuracy was 89.5%. For the visual attention task, we confirmed that the More/Less condition was less accurate than the All/Some condition (More/Less mean accuracy=85%. All/Some mean accuracy=95.8%. z=-3.82, p<.001).

For the sentence reading task, linear mixed modeling of raw reaction times for answering the questions did not yield residuals that were normally distributed, so we modeled log-transformed reaction times. Outliers were removed using median absolute deviation (outliers_mad function in ROutliers, version 0.0.0.3. Parameters b=1.4826, threshold=3 for normal distribution). Reaction times were slower for the Plausible than the Implausible condition (Mean log reaction time for Plausible=.37; Implausible=.29; t(41.1)=3.87, p<.001). For Long versus Short WM, the difference was not significant (Mean log reaction time for Long WM=.30; Short WM=.28; t(40.64)=0.7, p=.49). For control sentences, mean log reaction time was .34. For the visual attention task, we log-transformed the reaction times and removed outliers using the same procedure as above. Reaction times were slower for the More/Less than the All/Some condition (Mean log reaction time for More/Less=.88; All/Some=.65; t(37.29)=10.21, p<.001).

Overall, the norming study confirmed that the Plausible condition was more susceptible to lingering misinterpretation than the Implausible condition, yielding less accurate and slower responses to the questions. For the visual attention task, the results confirmed that the More/Less condition was more difficult than the All/Some condition, with less accurate and slower responses. We did not find significant effects for the Long vs Short WM contrast, suggesting that this manipulation was subtle and may potentially yield only weak brain activation differences.

### fMRI Study

#### Participants

For the fMRI experiment, we recruited participants through flyers and online postings on the George Washington University’s Psychology research credit portal. Thirty-two participants (8 Male, 24 Female, 18-28 years, Median age=19) participated. All were right-handed, native English speakers with normal or corrected to normal vision from the Washington, DC, area. They self-reported no history of neurological disorders or brain injury or use of neuropsychiatric medications. All participants underwent MRI safety screening and provided written informed consent under a protocol approved by the George Washington University Institutional Review Board. They received $25 or course credit for their participation. Three participants were excluded—one did not complete all the imaging runs, one had poor accuracy (<50% across all conditions), and one belatedly reported an ADHD diagnosis—leaving 29 participants in the final analyses.

#### Materials and Procedure

Before entering the scanner, participants were familiarized with the two tasks that they would be doing during the experiment. For the sentence reading task, participants first received 8 practice trials with feedback. They repeated this practice until they got at least 7 out of the 8 trials correct (i.e., answered the comprehension question correctly). Subsequently, they received 10 practice trials with no feedback, resembling the task structure inside the scanner. Practice sentences repeated during different phases of the practice but did not appear during the experiment. For the visual attention task, participants first received 8 practice trials with feedback. They repeated this practice until they got at least 7 out of the 8 trials correct. Subsequently, they received 4 practice trials with no feedback, resembling the task structure inside the scanner. The sentences used in the practice trials did not appear during the experiment.

Inside the scanner, participants completed two runs each of the sentence reading and visual attention tasks, respectively. The order of the runs was counterbalanced, with half the participants completing the sentence reading task during the first and the third runs and the visual attention task during the second and the fourth, and the other half doing the reverse. Stimuli were the same as those used in the norming study. All stimuli were presented using E-Prime 2.0.

##### Sentence Reading Task

During the sentence reading task, participants silently read whole sentences that were presented one at a time on a computer screen. After reading each sentence, they answered a comprehension question. All text appeared in black font on a white background. Responses were made using the index, middle, and ring fingers of the right hand on the left, middle, and right buttons, respectively. Participants were asked to press the middle button to indicate completion of reading. Subsequently, they answered the yes/no comprehension question using the left button for ‘Yes’ or the right button for ‘No’. Sentences were classified into five conditions, namely, Implausible, Control, Plausible, Long WM and Short WM (as in examples 1-5 above).

Each of two runs contained 105 sentences (21 in each condition) with their corresponding comprehension questions. For the sentences with subordinate clauses (Implausible, Plausible, Control), the comprehension question tested the thematic role assigned to the noun phrase following the verb. For garden-path Implausible and Plausible sentences, the correct answer was No (e.g., Sentence: *As the men wrestled the rivals arrived at the gymnasium;* Question: *Did the men wrestle the rivals?*). This is because the sentence structure indicates that “the rivals” are the subject of arriving and not the object of wrestling. The proportion of “Yes” responses is commonly used to infer the extent to which comprehenders hold on to or let go of the original misinterpretation (see e.g., Christianson et al., 2001). To partially balance out the type of correct response within a block, for non-garden-path Control sentences, the correct answer was Yes.^2^ In the working memory conditions, the comprehension question tested whether participants remembered various details from the content of the sentences. Responses within each condition (Long or Short WM) were split roughly evenly between Yes and No (11 vs. 10 out of 21). See Appendix for a full list of sentences and questions. Trials were pseudorandomly ordered such that the same condition did not occur more than 2 times in a row and the correct answer was not the same more than 3 times in a row.

For sentences with subordinate clauses, 42 verbs appeared once each in Implausible, Plausible and Control conditions, within the subordinate clause, across the whole experiment. If a verb appeared in the Implausible condition in the one run, it appeared in the Plausible condition in the other run and vice versa. A verb appeared in the Control condition in the same run as the Implausible or the Plausible condition roughly half the time. The main clauses of the sentences contained 94 different verbs that appeared between 1-3 times. Within each condition, half of the sentences began with “While” and half with “As” (21 each). Sentences in the three conditions had similar length (number of characters) (Implausible Mean=59.1, SD=4.7; Plausible Mean=57.6, SD=4.5; Control Mean=59, SD=7.6). Twenty-two proper names (e.g., Kendra, Sam) were used once each in each condition. Implausible and Plausible sentences were further matched such that they had the same subordinate clause verbs and had overlapping lexical frequencies of the subordinate clause nouns, main clause verbs and main clause nouns (Corpus of Contemporary American English (COCA), Davies 2008. Subordinate clause Noun: Implausible Mean=81327, SD=104375; Plausible Mean=73553, SD=169493. Main clause verb: Implausible Mean=79536, SD=105548; Plausible Mean=70444, SD=130756. Main clause Noun1: Implausible Mean=39292, SD=66760; Plausible Mean=55844, SD=91346. Main clause Noun2: Implausible Mean=91092, SD=157642; Plausible Mean=52769, SD=75203).^3^

For the working memory stimuli, the Long and Short WM conditions contained the same content with the two coordinated clauses in alternate orders (e.g., Long WM: *Josh went to see a foreign film and bought himself a variety of snacks*; Short WM: *Luke bought himself a variety of snacks and went to see a foreign film*). They did not differ in length (Mean number of characters: Long WM=88, Short WM=88.1).

Each sentence reading trial began with a 50 ms fixation cross followed by a jittered interval (min=1503, max=2996, mean=2256) and then the sentence, which was shown until the participant responded or for a maximum of 7 seconds. This was followed by a 50 ms fixation cross, another jittered interval (min=1510, max=2992, mean=2243), and then the comprehension question, which appeared until the participant responded or for a maximum of 4 seconds. See Figure 2a for an example trial.

**Figure 2:**
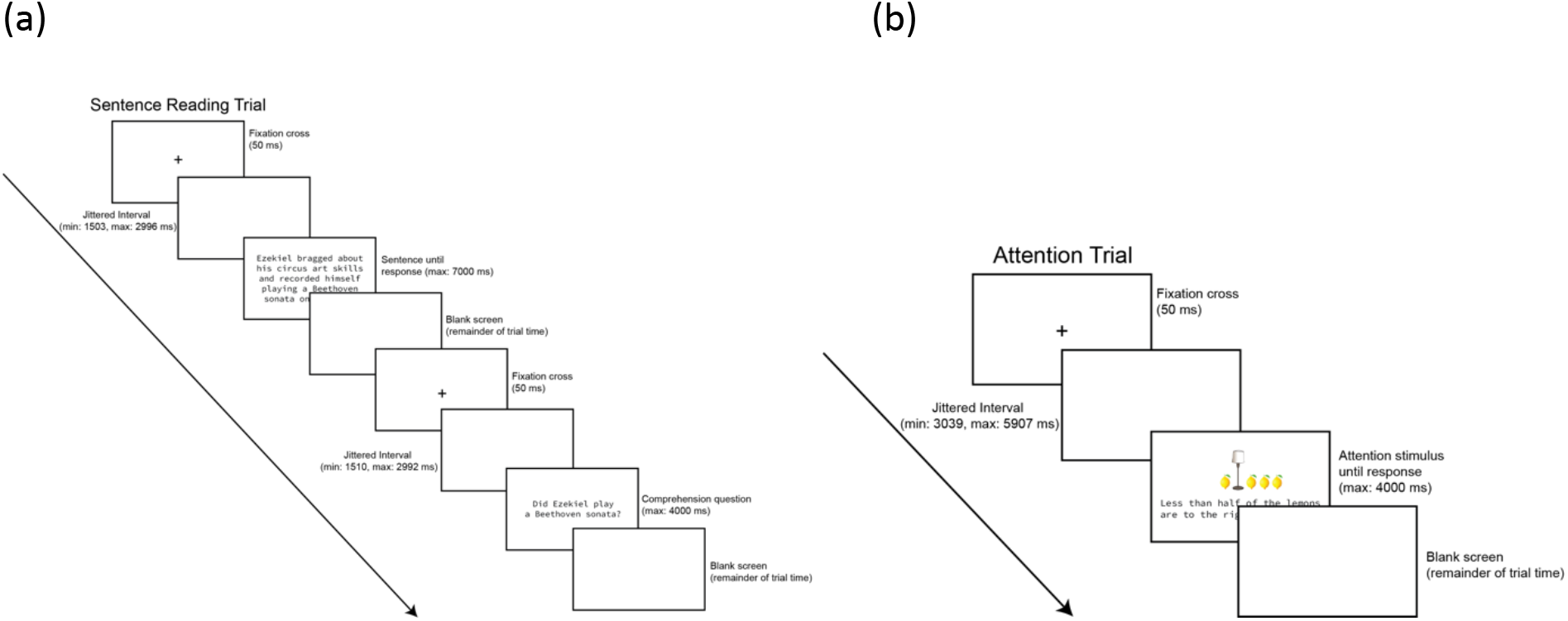
Trial structure in (a) the sentence reading task and (b) the visual attention task.

##### Visual Attention Task

In the visual attention task, participants saw a colored image and a sentence below it and were asked to evaluate whether the sentence described the image accurately. They responded using the index and ring fingers of their right hand on the left and right buttons to indicate Yes and No, respectively. Sentences were displayed in black text on a white background. The image featured a lamp in the center of the screen and various objects to the left and/or right of the lamp. Sentences were classified into two conditions: More/Less or All/Some (see examples 6-7 above).

Each participant completed 2 runs with 42 trials each (21 More/Less, 21 All/Some). Each quantifier appeared 10-11 times within a run. The correct response was Yes or No half of the time (21 each, split roughly evenly, 10-11 times in each condition). Across runs, each condition contained “right” versus “left” an equal number of times (21 each). Forty-two nouns appeared once each in the two conditions. See Appendix for a full list of sentences. Trials were pseudorandomly ordered such that the same quantifier did not occur more than 2 times in a row, nouns did not repeat in consecutive trials, and left/right and yes/no correct answer did not occur more than 3 times in a row.

Each trial began with a 50 ms fixation cross followed by a jitter (min=3039 ms, max=5907 ms, mean=4483 ms) and then the stimulus, which was shown until the participant responded or for a maximum of 4000 ms. See Figure 2b for an example trial.

#### fMRI Acquisition and Analyses

Structural (TR=1900 ms, TE=2.52 ms, Slice thickness=1 mm, 176 slices) and functional images (TR=1400 ms, TE=29 ms, Interleaved multi-slice mode, Slice thickness=2.3 mm, Flip angle=90°) were acquired using a 3T Prisma-Fit scanner. All scans took place at the Center for Functional and Molecular Imaging at Georgetown University. Analyses were conducted using FSL. Out-of-brain voxels were removed. Pre-processing included motion correction using MCFLIRT, interleaved slice timing correction, spatial smoothing (FWHM=5 mm), and high-pass filtering (100 s). Images were registered to the MNI 2-mm template. All statistical models included the standard motion parameters and motion outliers (fsl_motion_outliers). For the sentence reading task, the model included 10 events—two jitters, two fixations, the five conditions (Implausible, Control, Plausible, LongWM, ShortWM), and the question. For the visual attention task, the model included 4 events: jitter, fixation, and the two conditions (More/Less, All/Some). Jitters were of variable durations (see Figure 2) to facilitate separating activation for the sentence from activation for the question and/or separating activation for different trials.

In ROI analyses, we functionally profiled left hemisphere regions identified from the contrast of garden-path Implausible sentences versus non-garden-path Control sentences (Z threshold=3.1, Cluster p<.05), which could include regions involved in cognitive control, syntactic analysis, semantic/pragmatic processing, working memory, and general task demands. The large left temporal cluster from this analysis was split into an anterior temporal cluster (intersection with left anterior MTG from the Harvard-Oxford atlas) and a posterior temporal cluster (all other voxels posterior to the anterior part). The left frontal cluster from the analysis was split into a left frontal (language) cluster based on intersection with the language network, and a left frontal (language/MD) cluster based on intersection with both the language and the MD networks from the probabilistic functional atlas of Fedorenko and colleagues (Lipkin et al., 2022). Using these group-based clusters, we defined subject-specific ROIs as 5 mm spherical regions around the peak Implausible activation coordinate for each subject.^4^ Thus, there were 4 ROIs per subject—left posterior temporal, left anterior temporal, left frontal (language) and left frontal (language/MD). Within each ROI, Implausible will have higher activation than Control by definition because we identified voxels for analysis based on that criterion. Our main interest was in determining which of Plausible, Long WM, and Short WM also showed higher activation compared to Control. Accordingly, we conducted 3 paired t-tests contrasting each of those conditions with Control (Bonferroni correction for p<.05/3 = p<.0166) within each ROI. Additionally, to determine if the regions showing increased recruitment for one or more of the three sentence types also showed the signature of increased activation based on attention demands, we examined the contrast of More/Less versus All/Some from the visual attention task within each ROI. Together, these analyses were used to functionally profile different regions based on the logic outlined in Table 1.

In addition to the ROI analyses, we conducted whole-brain analyses contrasting closely matched conditions for each type of stimulus: (i) Implausible vs Plausible garden-path; (ii) Long vs Short WM; and (iii) More/Less vs All/Some visual attention. This was intended to provide supplementary information about other potential regions engaged for comprehending sentences that were difficult along different dimensions (semantic/pragmatic processing, working memory, visual attention). In each case, we also examined the reverse contrast (e.g., Short vs Long WM) for completeness.

## Results

### Behavioral Results: Sentence Reading Task

For the sentence reading task, we analyzed reading time for the sentence, and accuracy and reaction time for responding to the comprehension question. For reading times, trials where the participant did not respond within the time allowed were replaced with the maximum duration (7000 ms). Residuals from modeling raw reading times were normally distributed, so we did not log-transform them. Reading times were adjusted for length. There were no outliers. Mean length-adjusted reading time for the different conditions were as follows: Control=43.9, Implausible=41.6, Plausible=38.2, Long WM=-30.7, Short WM=-92.9. Compared to Control, Implausible and Plausible sentences were not significantly different (p’s>.9). Short WM had a significantly shorter length-adjusted reading time (t(2406.00)=-2.93, p<.01). For Long WM, the effect was in the same direction but not statistically significant (t(2406.00)=-1.52, p=.13). Thus, the working memory stimuli, although longer, were read at a quicker rate, suggesting that these sentences with coordinate clauses were easier to read than the sentences with subordinate clauses (Control, Implausible, and Plausible).

For comprehension questions during the reading task, mean accuracies in the different conditions were: Control=88.5%, Implausible=89.8%, Plausible=78.2%, Long WM=82.8%, Short WM=89%. All conditions were above chance (exact binomial test p’s<.001), indicating that participants attended to the task. We compared closely matched conditions that had similar questions (Implausible vs Plausible and Long WM vs Short WM). The Plausible condition had significantly lower accuracy than the Implausible condition (z=-8.40, p<.001), consistent with our predictions and previous findings on lingering misinterpretations. The Long WM condition had significantly lower accuracy than the Short WM condition (z=-4.78, p<.001), consistent with predictions based on working memory demands. For reaction times, trials where the participant did not respond within the time allowed were replaced with the maximum duration (4000 ms). We log-transformed the reaction times and removed outliers (using the same procedure as for the Norming study). Mean log reaction times for the different conditions were as follows: Control=7.03, Implausible=6.94, Plausible=7.0, Long WM=7.29, Short WM=7.30.

Comparing Implausible and Plausible conditions, the latter had a significantly longer reaction time than the former (t(1980)=3.58, p<.001). Long and short WM conditions did not differ from one another (p>.6). To summarize, participants showed lower accuracy and longer reaction time to questions about the ambiguous portion of Plausible than Implausible sentences, indicating more lingering misinterpretations for the former that cannot be attributed to a speed-accuracy tradeoff. For working memory, participants showed lower accuracy and equivalent reaction time to questions about Long than Short WM sentences, tentatively indicating some difficulty with the former sentence type.

### Behavioral Results: Visual Attention Task

For the visual attention task, we analyzed accuracy and reaction time. Mean accuracies for the More/Less and All/Some conditions were 80.5% and 92.1%, respectively. Both were above chance (binomial p’s<.001). Comparing the two conditions, participants were significantly less accurate on More/Less (z=-8.12, p<.001), consistent with our predictions. Reaction times were log-transformed. There were no outliers identified using the same procedure as above. Log-transformed reaction times for the two conditions were as follows: More/Less=7.75, All/Some=7.54. There was a significant effect of condition (t(2050)=20, p<.001), with responses being slower in the More/Less condition. Together, these analyses confirmed that More/Less taxed visual attention more than All/Some, as intended, resulting in lower accuracy and longer reaction time.

### fMRI Results: Sentence Reading Task

Garden-path Implausible sentences showed more activation over non-garden-path Control sentences in left temporal and frontal areas that fell within Fedorenko and colleagues’ language as well as MD networks (Figure 3. See SI #1 for the list of clusters). Subject-specific ROIs were defined around the peak activation for Implausible sentences for each subject within these clusters (see insets in Figure 4).

**Figure 3:**
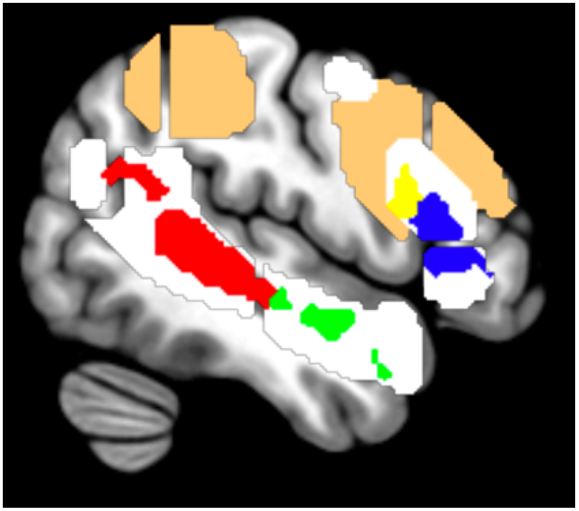
Implausible versus Control contrast revealed activation in left posterior temporal (red), left anterior temporal (green), left frontal (language) (blue), and left frontal (language and MD) (yellow) regions. Clusters are superimposed on Fedorenko and colleagues’ language (white) and MD (tan) networks.

**Figure 4:**
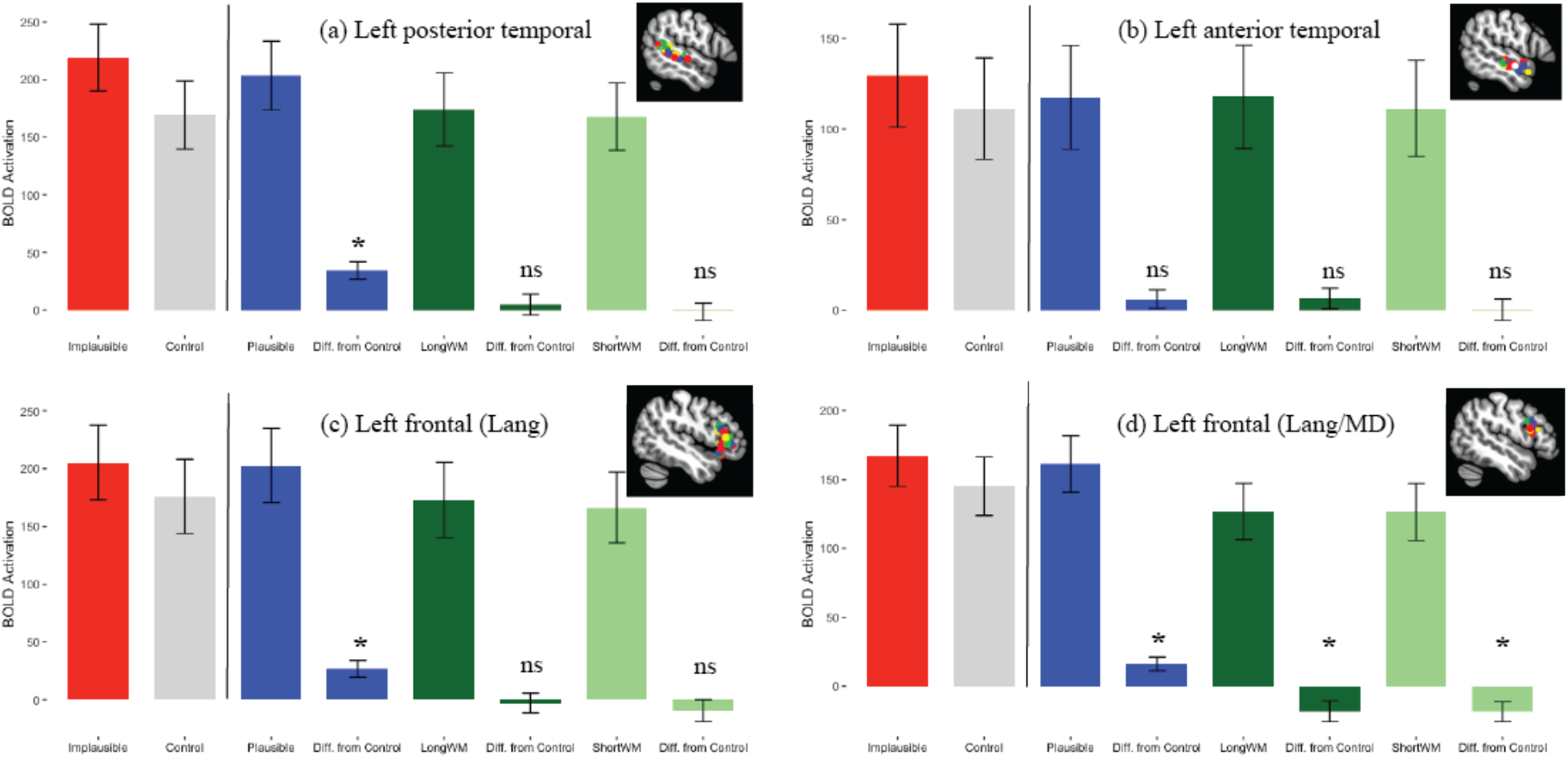
Results for the sentence reading task in (a) Left posterior temporal (b) Left anterior temporal (c) Left frontal (Language) and (d) Left frontal (Language/MD) ROIs. ROIs were obtained from the contrast of Implausible > Control, so only information to the right of the vertical line in each panel should be interpreted. *=significant at Bonferroni-corrected threshold, ns=not significant for the difference (Diff.) between each condition and control. Brain image inset in each panel shows the spherical ROIs for different participants.

Figure 4 shows activation for the different conditions in the sentence reading task within the left posterior temporal, left anterior temporal, left frontal (language), and left frontal (language/MD) ROIs. Paired-sample t-tests revealed that in the left posterior temporal ROI, Plausible showed increased activation over Control (t(28)=4.81, p<.001. Figure 4a). Here and elsewhere, means and standard errors are shown per condition for information, but the bars marked “Diff.” are the ones to interpret corresponding to the inferential statistics. Long and Short WM conditions did not show a significant difference from Control (Long: t(28)=.57, p=.58; Short: t(28)=-.22, p=.82). The same pattern was observed in the left frontal (language) ROI (Figure 4c. Plausible: t(28)=3.97, p<.001; Long WM: t(28)=-.36, p=.72; Short WM: t(28)=-1.07, p=.30). In the left anterior temporal ROI, none of the three comparisons revealed significant effects (Figure 4b. All p’s>.2). Finally, in the left frontal (language/MD) ROI (Figure 4d), Plausible showed increased activation compared to Control (t(28)=3.50, p=0.002), while Long and Short WM showed decreased activation (Long: t(28)=-2.58, p=.015, Short: t(28)=-2.77, p=.001). To summarize, a pattern consistent with involvement in cognitive control or syntactic reanalysis was found in the left frontal (language) and the left posterior temporal ROIs, which showed increased activation for Plausible over Control (in addition to Implausible over Control). The left frontal (language/MD) ROI was subtly but importantly different from the left frontal (language) ROI in how it responded to the Long and Short WM conditions (see more in Discussion). The left anterior temporal ROI was the only ROI that was activated for the Implausible but not the Plausible condition relative to Control (see more below).

Whole-brain analysis for Implausible versus Plausible revealed a significant effect in left anterior temporal cortex only (Figure 5), consistent with the ROI results. For Long versus Short WM, there were no significant clusters. In both cases, the reverse contrast (Plausible > Implausible, Short WM > Long WM) revealed no significant effects.

**Figure 5:**
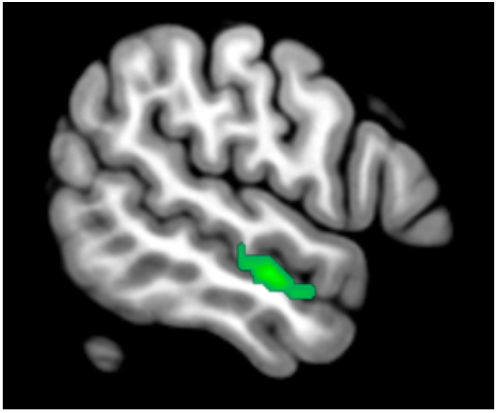
Whole-brain analysis results Implausible vs. Plausible sentences. Single cluster. Peak coordinate at MNI (−56, -12, -14)

Because the planned ROI and whole-brain analyses did not identify any areas sensitive to working memory demands, we conducted a post-hoc whole-brain analysis to see if Long WM showed more activation than Control in any brain regions. This revealed several occipital, temporal and medial frontal clusters (see SI #2), suggesting that we could detect activation related to the WM manipulation and that the null results in the left frontal cortex were not solely due to low power. In the left perisylvian cortex, there were two temporal clusters. The pattern of activation within the larger of these clusters is shown in Figure 6. As before, we defined spherical ROIs around subject-specific peaks within the cluster. Long WM showed more activation over the Short WM as well as the Implausible condition (Long WM vs Short WM: t(28)=2.12, p<.05; Long WM vs Implausible: t(28)=2.15, p<.05). These results should be interpreted with caution because the analysis was post-hoc and we looked within ROIs defined for showing high Long WM activation. However, they provide additional context for interpreting the patterns shown in Figure 4 (see Discussion).

**Figure 6.**
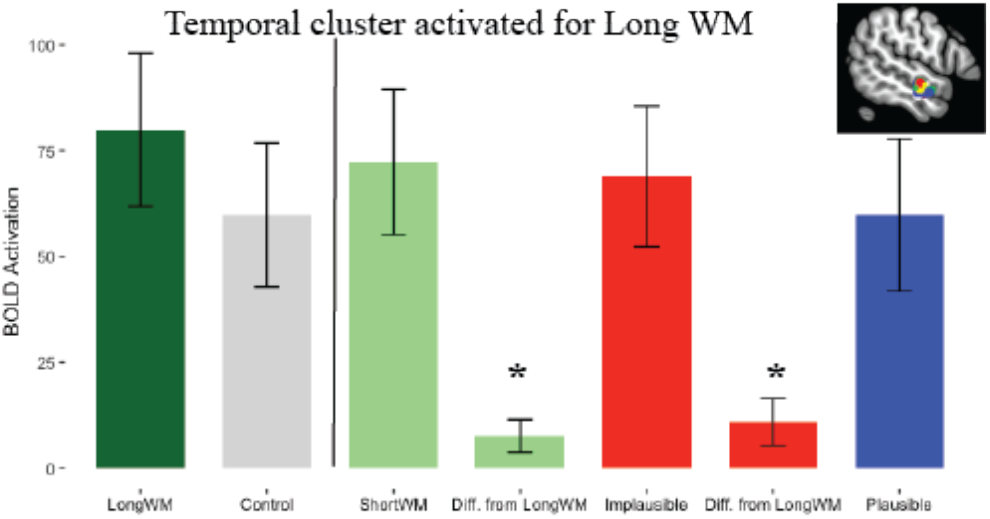
Results for the sentence reading task in subject-specific left temporal ROIs obtained from the post-hoc contrast of LongWM > Control. Only information to the right of the vertical line in each panel should be interpreted. *=p<.05.

### fMRI Results: Visual Attention Task

Whole-brain analysis of More/Less versus All/Some found a significant effect in the visual cortex, consistent with the hypothesized need for greater visual processing in this condition (Figure 7a). The reverse contrast (All/Some versus More/Less) revealed significant effects in several bilateral anterior and posterior regions (Figure 7b), indicating that All/Some recruited additional—including linguistic—processing that was different from the visual processing used for More/Less.

**Figure 7.**
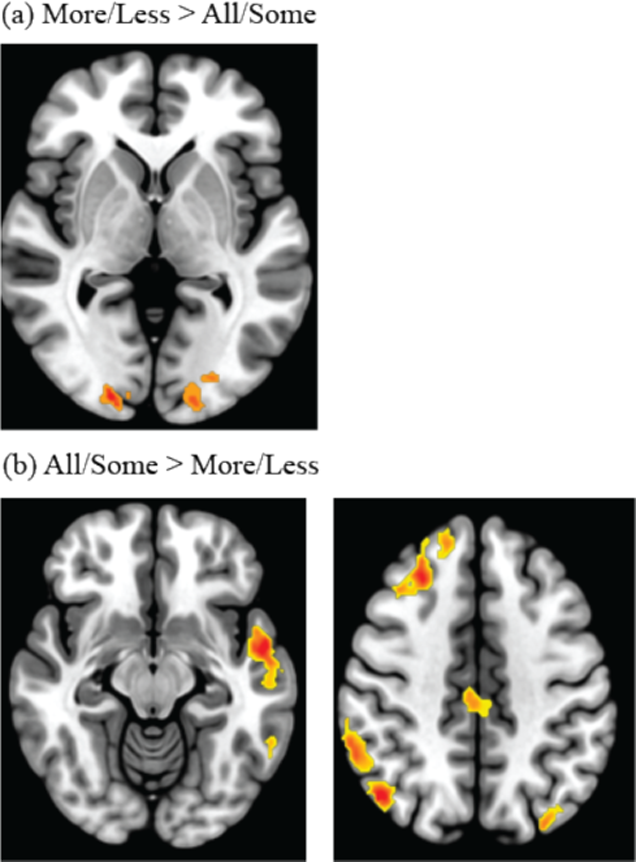
Whole-brain analysis results for the visual attention task

Looking in the ROIs examined for the sentence reading task (Figure 8), All/Some showed more activation than More/Less in both temporal regions (left posterior temporal: t(28)=3.72, p<.001; left anterior temporal: t(28)=2.28, p=0.03). The difference was marginally significant in the left frontal (language) ROI (t(28)=1.87, p=0.07) and not significant in the left frontal (language/MD) ROI (t(28)=0.28, p=0.78). Thus, the results suggested that All/Some recruited more linguistic processing than More/Less in temporal regions. Similar to the results for the sentence reading task, the two frontal ROIs were subtly but importantly different, suggesting a gradient for language-specificity (see Discussion).

**Figure 8.**
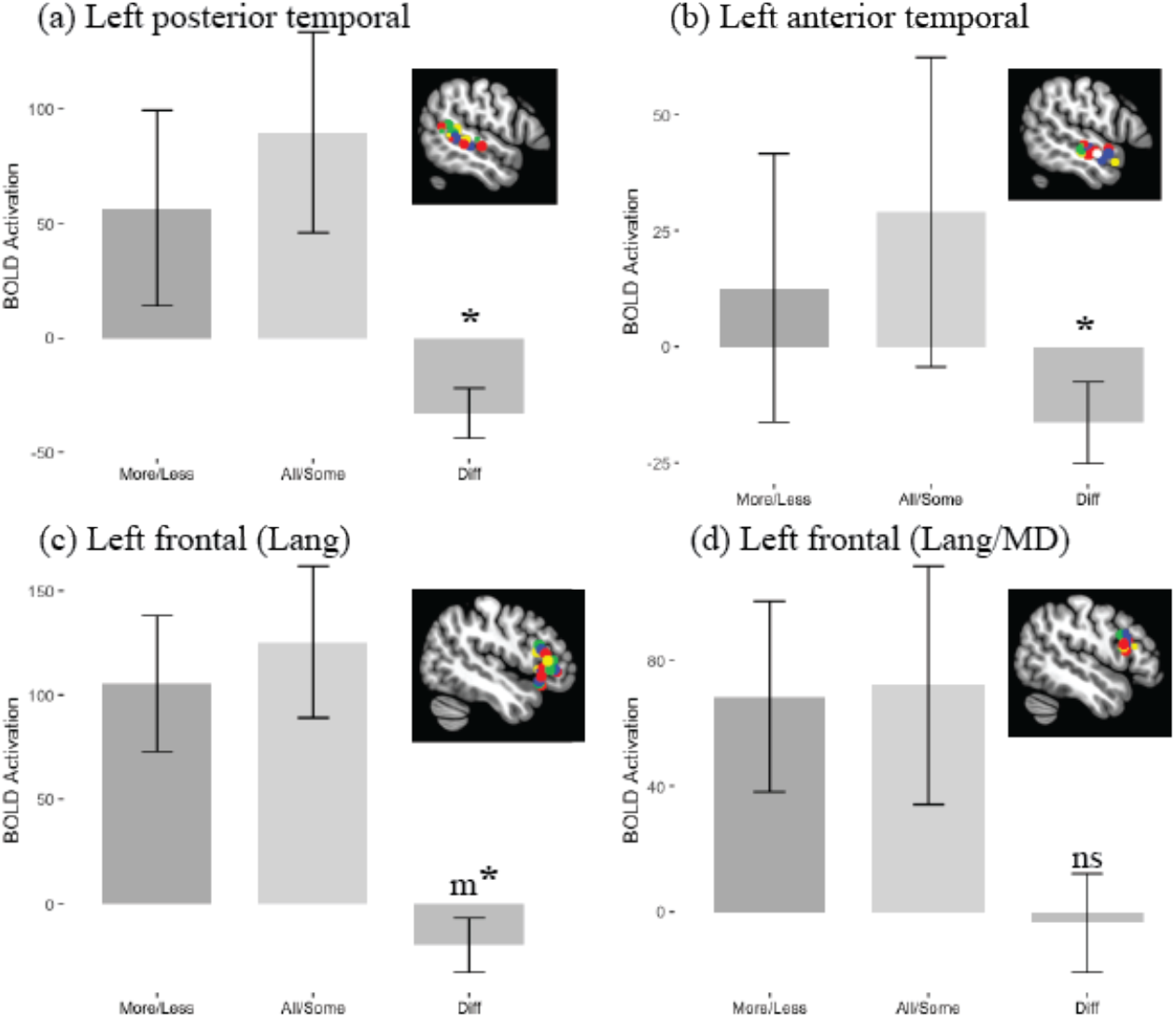
Results for the visual attention task in (a) Left posterior temporal (b) Left anterior temporal (c) Left frontal (Language) and (d) Left frontal (Language/MD) ROIs (same as in Figure 4). *=significant, m*=marginally significant, ns=not significant for the difference (Diff.) between conditions.

## Discussion

The goal of this study was to fractionate difficulty during sentence comprehension. Difficult-to-comprehend sentences are often difficult in more than one way. Therefore, we used a functional profiling approach to isolate different processes that can contribute to the interpretation of such sentences. Our results suggest that sentence comprehension recruits regions that lie on a spectrum of specificity to generality with respect to the representations and processes that are involved. On the highly specific end, hard-to-understand garden-path sentences recruited left temporal regions that are widely thought to support language-selective processing of linguistic representations—left posterior temporal cortex for syntactic analysis and left anterior temporal cortex for deeper semantic/pragmatic processing. On the highly general end, these sentences recruited dorsal left frontal cortex, which is linked to domain-general processing of a variety of representations (e.g., linguistic, spatial, mathematical) as part of the MD network. In between these extremes, these sentences also recruited ventral left frontal cortex, whose role is debated. This region could be involved in language-selective processing of linguistic representations (akin to the left temporal cortex. See e.g., Shain et al., 2022), a computationally general process of integration or cognitive control operating over linguistic representations (e.g., Hagoort, 2005; Thothathiri et al., 2012c), or a computationally general process of cognitive control operating over a variety of representations (e.g., Hsu et al., 2017). Below, we discuss each region in turn.

Multiple methodologies have converged on the left posterior temporal cortex as a possible neural correlate of syntactic processing. Damage to this region impacts comprehension, especially when sentences have to be interpreted using syntax and cannot be understood using semantic cues alone. Neuroimaging studies in neurotypical adults routinely report activation in this region when contrasting sentences that differ in their syntactic processing demands. In the present study, increased activation of this region for both types of garden-path sentences (Implausible and Plausible) compared to non-garden-path Control sentences is consistent with prior literature. Because these sentences tend to be misinterpreted initially, they have to be re-analyzed, requiring the recruitment of additional syntactic processing.

Neuropsychological evidence from semantic dementia and neuroimaging evidence from neurotypical individuals suggest that the left anterior temporal cortex is important for semantic processing. In the present case, it is notable that this was the only region that was recruited additionally for Implausible over Plausible garden-path sentences in the ROI analyses, out of the two temporal and two frontal ROIs. The whole-brain contrast of Implausible > Plausible also identified only this region. Prior psycholinguistic evidence suggests that Implausible and Plausible sentences both lead to garden-pathing and re-analysis but that the former induces *some* additional processing that enables the comprehender to successfully let go of the original misinterpretation (as confirmed by our behavioral accuracy results). Our ROI and whole-brain results juxtaposed with this evidence suggests that the additional processing might be semantic/pragmatic in nature, supported by the left anterior temporal cortex, potentially for assessing the compatibility of the original and the later interpretations of the sentence given our knowledge of the world (e.g., Can the wrestlers be wrestling the rivals if the rivals just arrived?). However, post-hoc analysis of LongWM versus Control sentences identified an ROI that was quite close to this region (compare the brain insets in Figure 4b and Figure 6). As designed, the LongWM sentences did not contain syntactic ambiguity and were not subject to any misinterpretation that should require deeper semantic/pragmatic processing. The observed proximity between these ROIs was therefore unexpected and we can only speculate about possible explanations. One possibility is that there is sub-division within the temporal cortex such that the anterior-most parts are engaged in semantic/pragmatic processing (for Implausible sentences, Figure 4b) and the parts that are more towards the middle are engaged in some process (e.g., verbal working memory) used for LongWM sentences (Figure 6). Alternatively, it could be that semantic/pragmatic processing consists of different sub-components, some of which were tapped more by our Implausible stimuli and others unexpectedly by our LongWM stimuli.

On the domain-general end of the spectrum, dorsal left frontal cortex (the left frontal (language/MD) ROI) revealed a profile consistent with general task-difficulty-based activation, as suggested by the previous literature on the MD network. We observed maximal activation in this region for the difficult sentences containing ambiguity/conflict, namely Implausible and Plausible. But we also saw more activation here for Control over LongWM and ShortWM. Thus, increased activation of this ROI was not linked to ambiguity/conflict alone but seemed to track general difficulty. Sentences with the highest length-adjusted reading times (Implausible, Plausible, Control) showed the most activation. Further, this ROI differed from the other three ROIs that were linked to the language network. Left posterior temporal, left anterior temporal, and left frontal (language) regions all showed a pattern of All/Some > More/Less in the visual attention task. This suggests that even though More/Less was harder as measured via accuracy and reaction times and the need for greater visual attention, All/Some imposed higher linguistic demands. By contrast, the left frontal (language/MD) ROI, while proximal to the left frontal (language) ROI, showed equivalent activation between More/Less and All/Some, suggesting that it might be sensitive to difficulty in a more general way that encompasses both linguistic and non-linguistic task demands. This distinction between ventral and dorsal regions within the left frontal cortex is consistent with prior evidence that suggests more domain-generality as we move more dorsally (Hagoort, 2005; Fedorenko et al., 2013).

Last but not least, we observed increased activation for Implausible and Plausible garden-path sentences over Control sentences within ventral left frontal cortex (the left frontal (language) ROI), consistent with a role for this region especially in cases of ambiguity and conflict. This is consistent with either cognitive control or syntactic processing (see Table 1). Hsu et al. (2017) demonstrated that several conflict resolution tasks that do not involve syntax (Stroop, N-back with lures, and Recent Negatives) showed activation within the same subject-specific voxels within ventral left frontal cortex as a syntactic ambiguity task. This suggests that this region is involved in a computationally general process, namely cognitive control, that can resolve conflict during different tasks (not just sentence processing). The authors suggested that functional specificity for particular representations might arise at the network level, wherein ventral left frontal cortex interacts with different brain regions for different kinds of representations. By comparison, Fedorenko and colleagues have argued that ventral frontal (and other) regions within their language-selective network are selectively activated for linguistic tasks. At the representational level, this can be successfully reconciled with Hsu et al. (2017) because all tasks in that study involved some kind of linguistic representation (letters, words, sentences). The results for the present study too are consistent with language-selectivity in this sense because the All/Some condition showed more activation than the More/Less condition in the ventral left frontal ROI, similar to the pattern in the left temporal ROIs (which uncontroversially deal with linguistic representations) and distinct from the pattern in the dorsal left frontal ROI (which is linked to the domain-general MD network). The question remains however about the type of *process* that is supported by this region. While Fedorenko and colleagues have suggested, either implicitly or explicitly, that this region is engaged in a language-specific process, others have pointed to co-localization among very different tasks (e.g., Stroop, sentence comprehension) to argue that the process is a more general one like integration or conflict resolution (Hagoort, 2005; Hsu et al., 2017; Thothathiri et al., 2012c). The present results are consistent with both the “more general” and the “more specific” perspectives. They show increased recruitment especially for sentences containing conflict, consistent with a general process of cognitive control within this region. However, we cannot rule out the possibility that this increased recruitment reflects something more language-specific like syntactic processing.

Within the left frontal ROIs, including left frontal (language), we did not find increased activation for Long WM over Short WM or Control sentences. Shain et al. (2022) reported that activation within the language-selective network, including left frontal cortical regions, corresponded with syntactic working memory demands. Our results do not fully accord with this conclusion. We observed increased activation for the hardest WM condition in other regions but not the left frontal cortex. However, the behavioral results and the lack of significant clusters for LongWM versus ShortWM suggest that our WM manipulation might have been weak. Therefore, this evidence should be interpreted cautiously.

The issue of difficulty during sentence comprehension and its relationship to cognitive control has come to the forefront in the past two decades of neuroimaging research because of growing evidence on two different fronts. On the one hand, a number of studies have now shown that frontal regions engaged during cognitive control tasks are also engaged during sentence comprehension and more causally, that triggering cognitive control facilitates the comprehension of sentences containing conflict. However, these demonstrations have raised questions about whether the evidence indicates a role for conflict resolution or cognitive control specifically, or whether the evidence can be interpreted more broadly in terms of “difficult” situations and the brain regions that support difficulty. Thus, whether cognitive control, in particular, is useful for sentence comprehension has been called into question. On a different front, other evidence has indicated a distinction between brain networks that support language processing and those that support broader demands like general-purpose difficulty. This evidence is usually interpreted as showing that language processing is mostly supported by language-selective processes and that any difficulty-related processing occurs in distinct brain areas (the MD network) and is only used under some circumstances. Thus, this strand of research too has questioned the relevance of cognitive control for sentence comprehension, relegating it to a cluster of general-purpose difficulty processing within the MD network.

Hsu et al. (2017) addressed this concern by examining whether the contrast between conflict and no-conflict trials in different tasks (Stroop, syntactic ambiguity, etc.) activated ventral left frontal cortex or the MD network. They found that 4 different conflict tasks converged in activating the same voxels within ventral left frontal cortex, supporting a role for this region in conflict resolution. In contrast, they did not observe such co-localization within the MD network. Therefore, the authors argued that general-purpose difficulty cannot account for their findings. The present study offers new complementary evidence on this debate. Like Hsu and colleagues, we found increased activation within ventral left frontal cortex for conflict versus no-conflict sentences. We also found activation for such sentences within more dorsal MD-related parts of the left frontal cortex, consistent with other researchers’ claims about general-purpose task demands and difficulty. However, through examining different kinds of sentences within the same participants, our results demonstrate the distinct functional profiles of ventral versus dorsal left frontal cortex. Both regions are engaged for sentences containing ambiguity and conflict but they show different recruitment patterns for other kinds of sentences that did not contain conflict and varied in difficulty. Thus, our results suggest that conflict resolution and the handling of general difficulty are not the same (recruiting different regions) and that they are both relevant during the comprehension of garden-path sentences.

More broadly, the dichotomization of language-selective versus MD regions in recent literature can be a useful empirical tool (as the present study too demonstrates), but we would suggest that this should not obscure the fact that the human brain can use multiple tools at its disposal to handle any given instance of sentence comprehension, be they domain-general or language-specific. Put another way, when a listener or reader encounters hiccups during sentence comprehension, both general-purpose processes that enable adjustment broadly to a difficult situation and more specific processes that adapt in a language-specific way can be helpful (see Sharer & Thothathiri, 2020 for additional evidence and discussion).

To summarize, we found that a variety of brain regions support comprehension when sentences are difficult to understand. Specifically, regions linked to syntactic analysis (left posterior temporal cortex), semantic/pragmatic processing (left anterior temporal cortex), linguistic cognitive control (ventral left frontal cortex), and general task demands (dorsal left frontal cortex) were all recruited when reading sentences that trigger conflicting interpretations that must be reconciled for accurate comprehension. Thus, difficulty during sentence comprehension is multi-faceted and different demands are handled by different brain regions.

## Appendix

### CCWM Stimuli

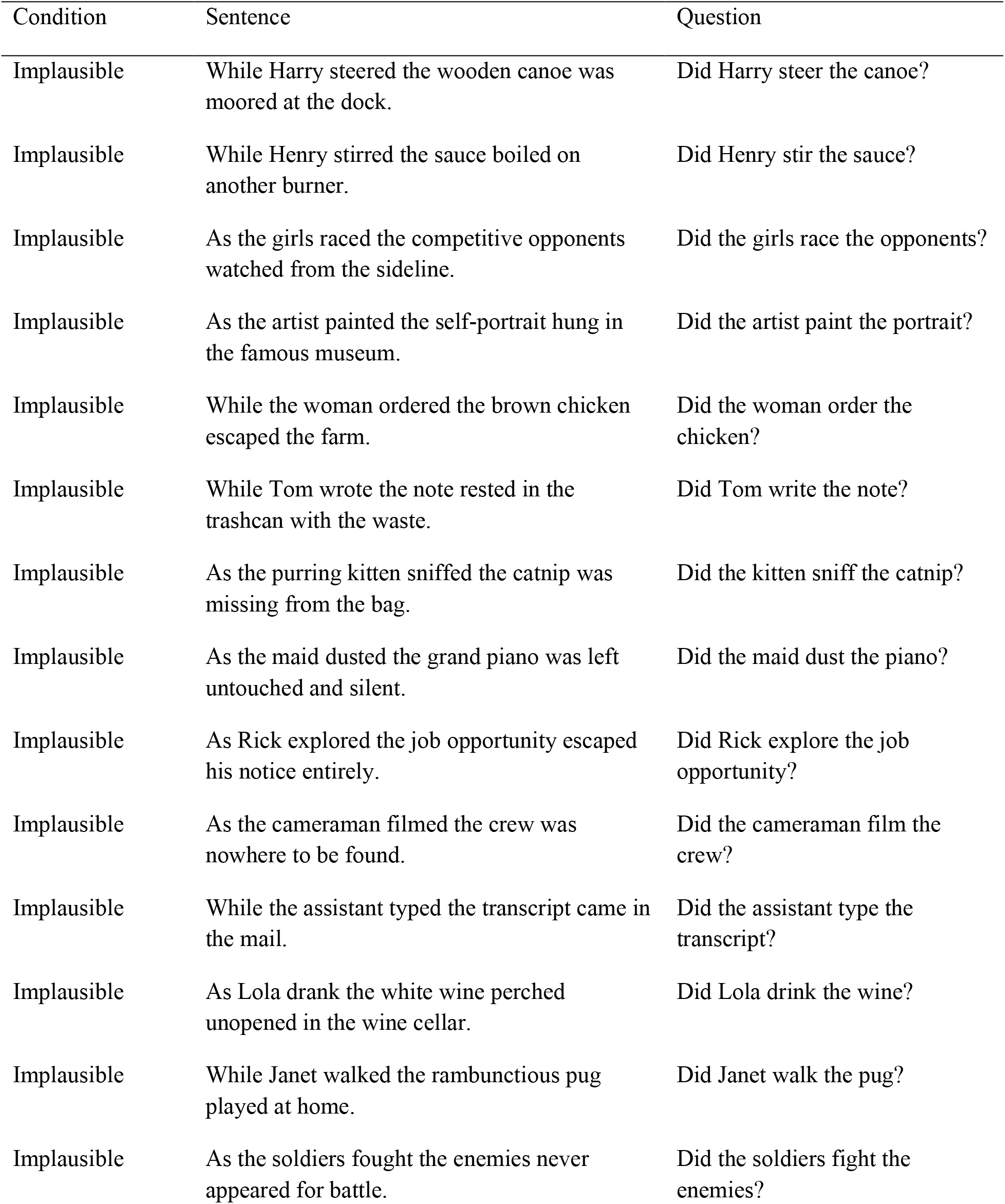
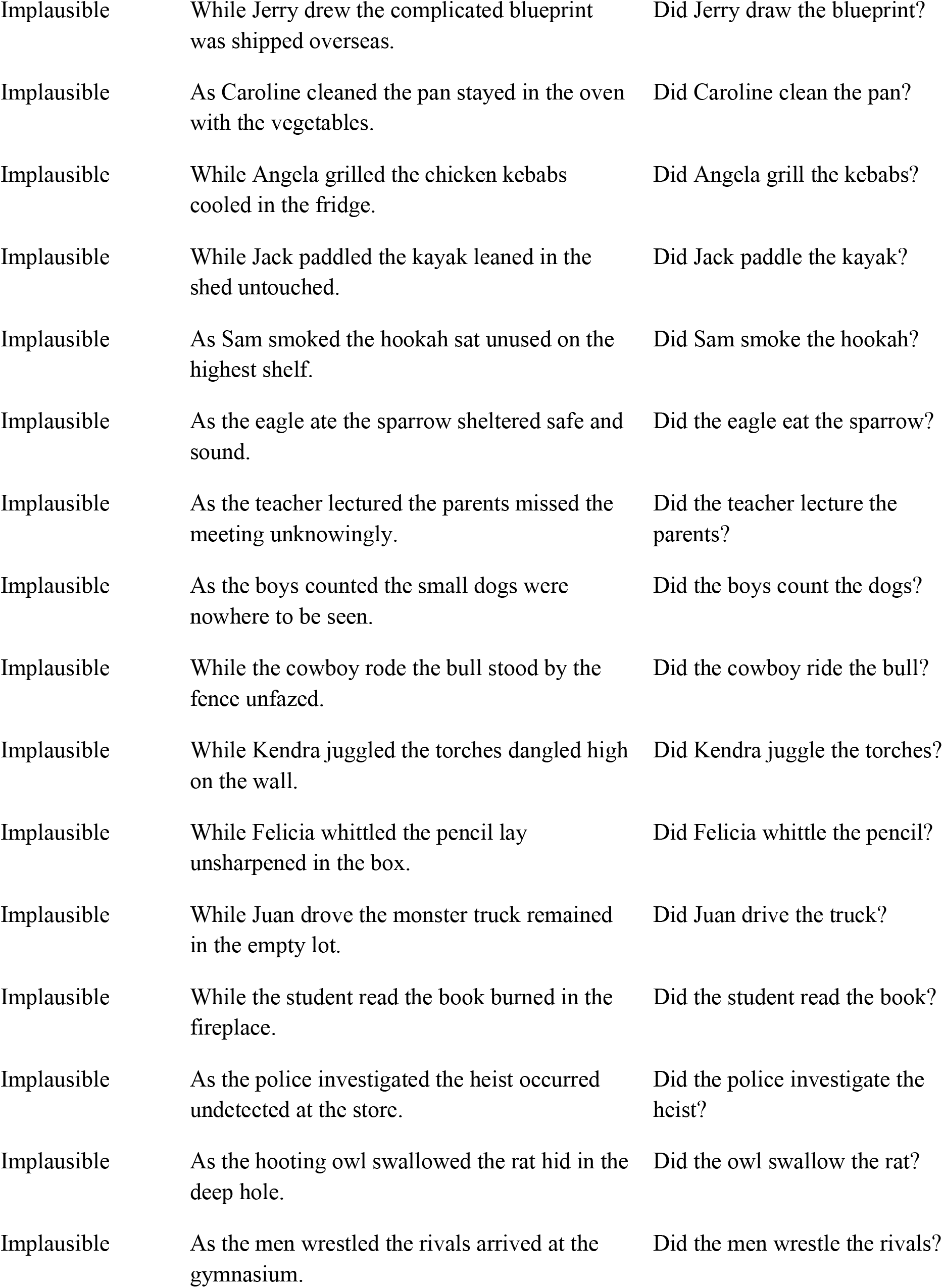
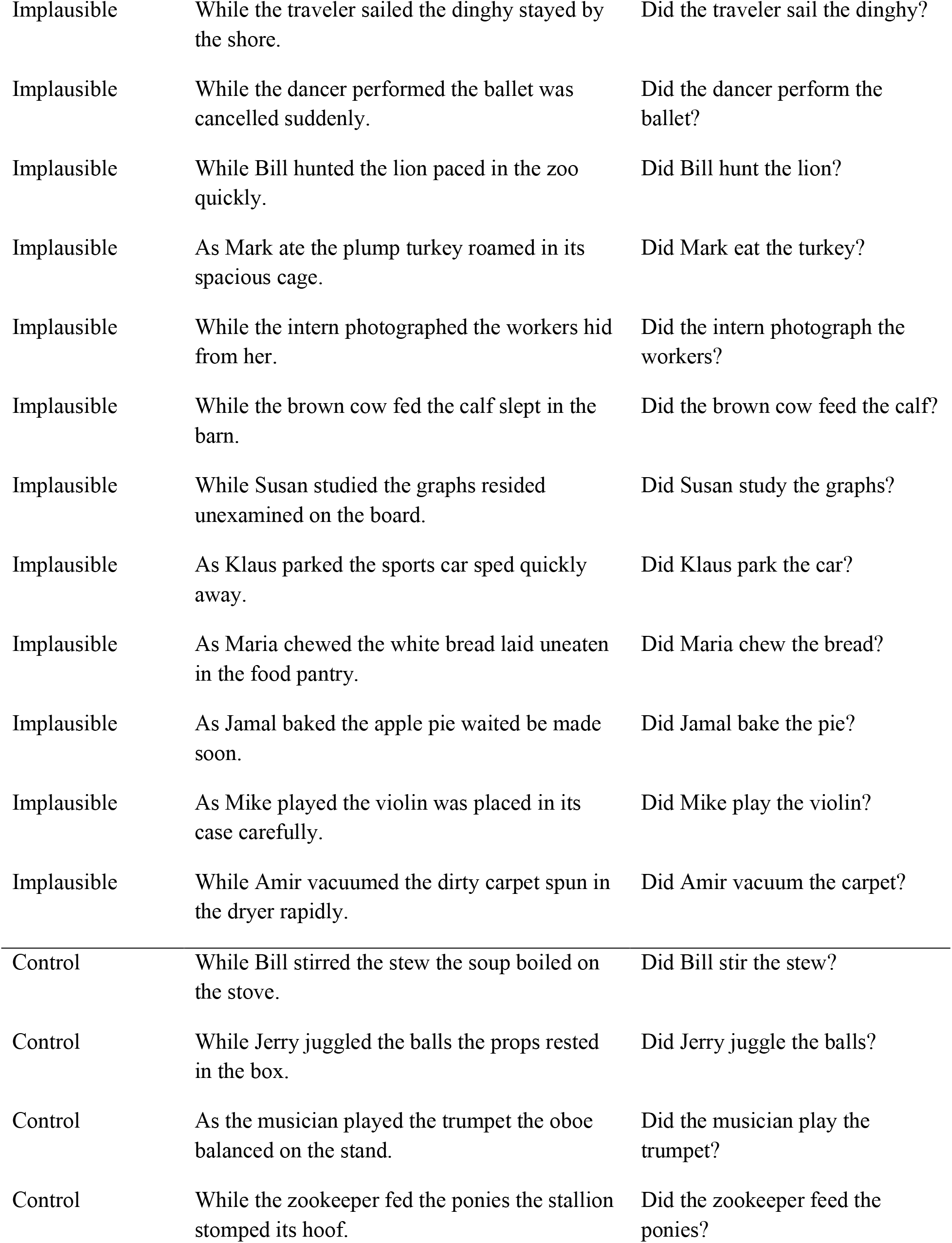
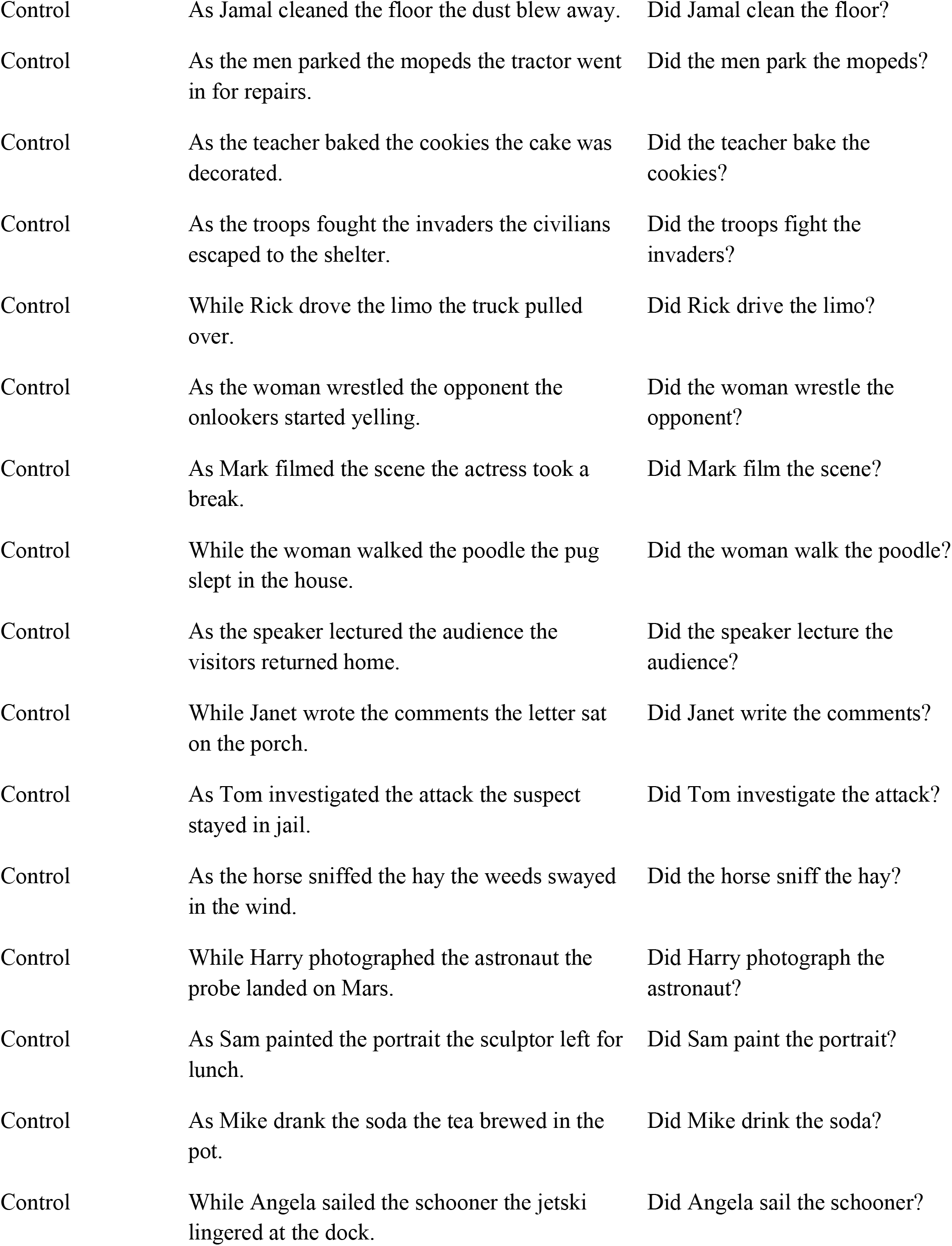
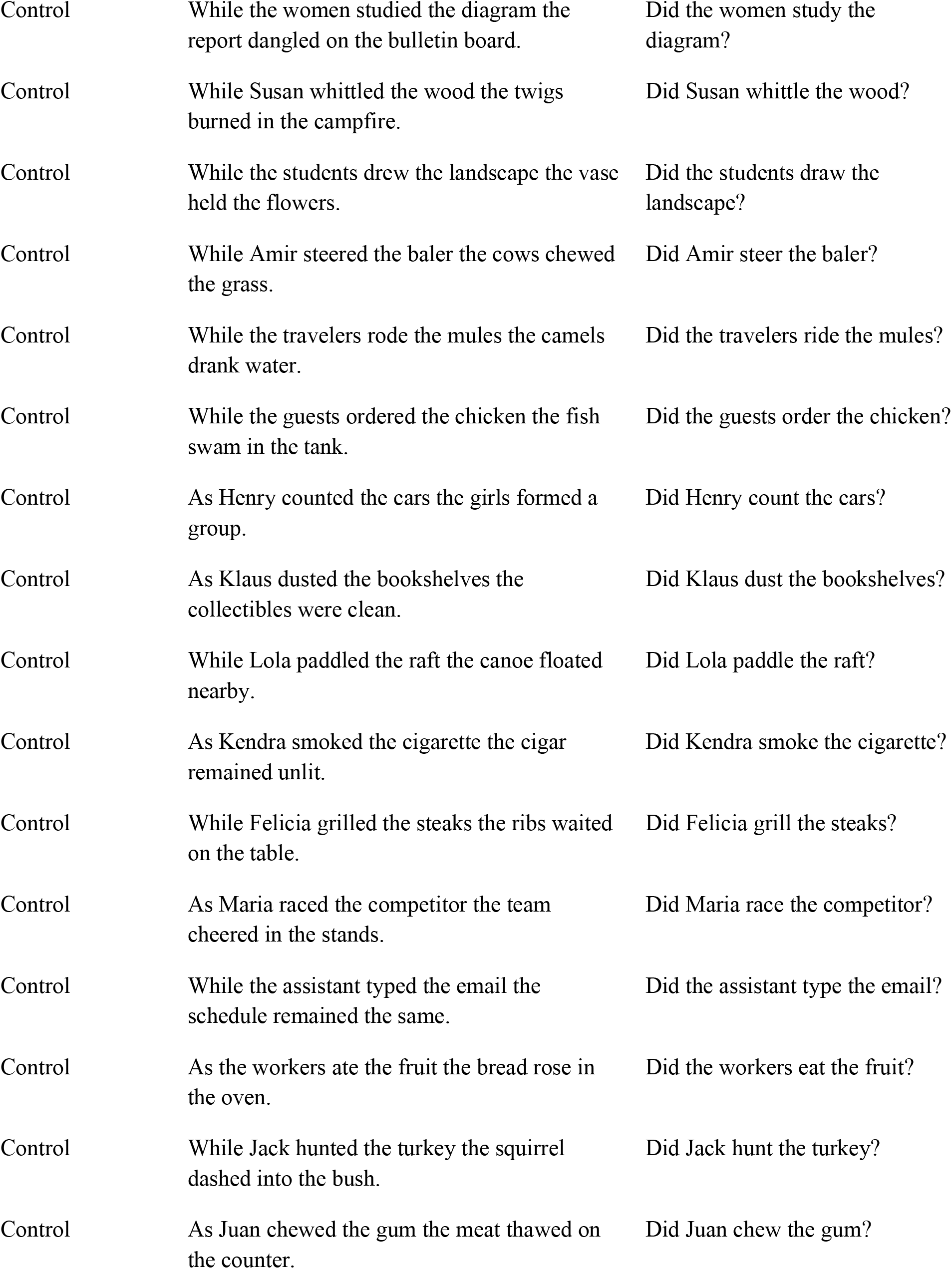
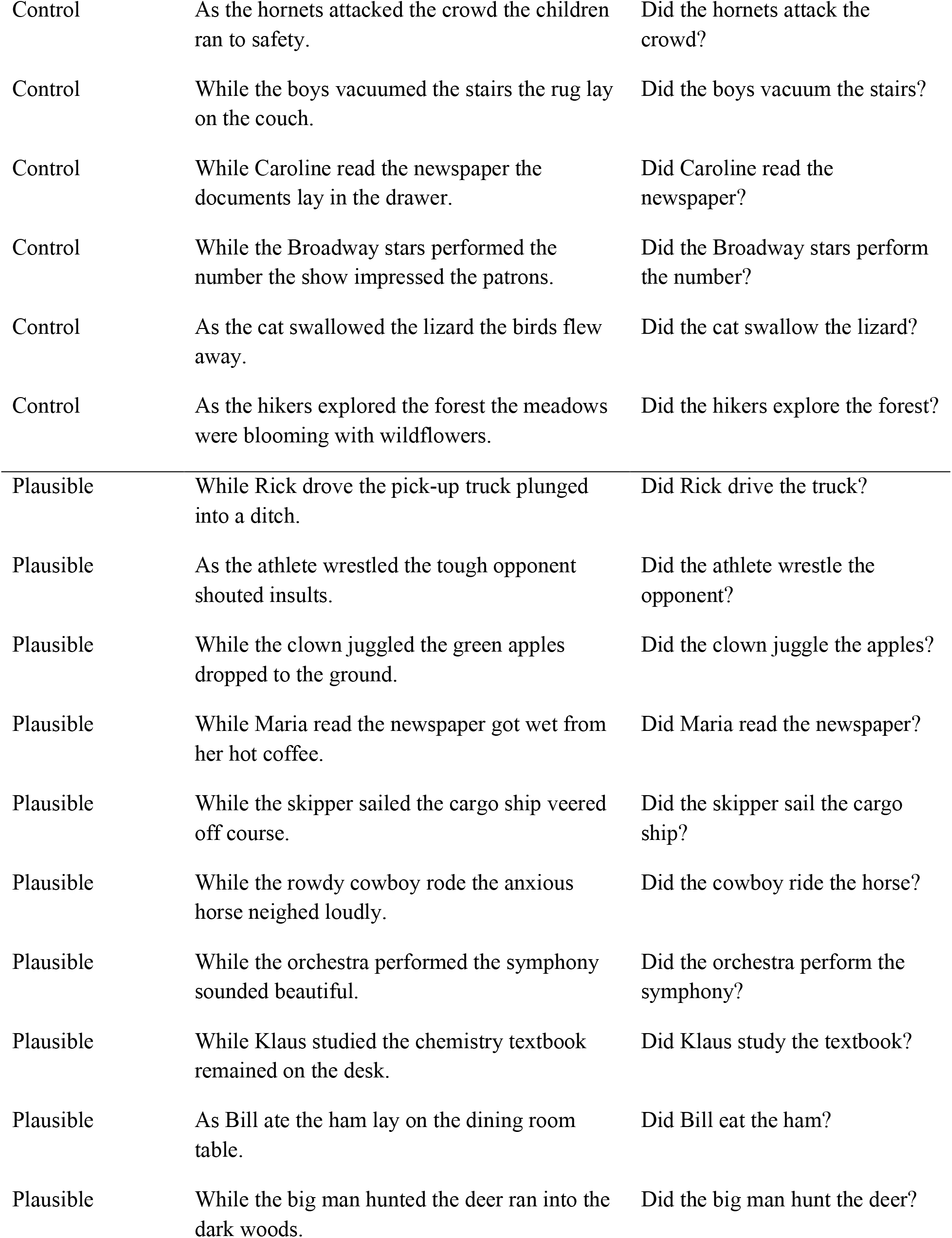
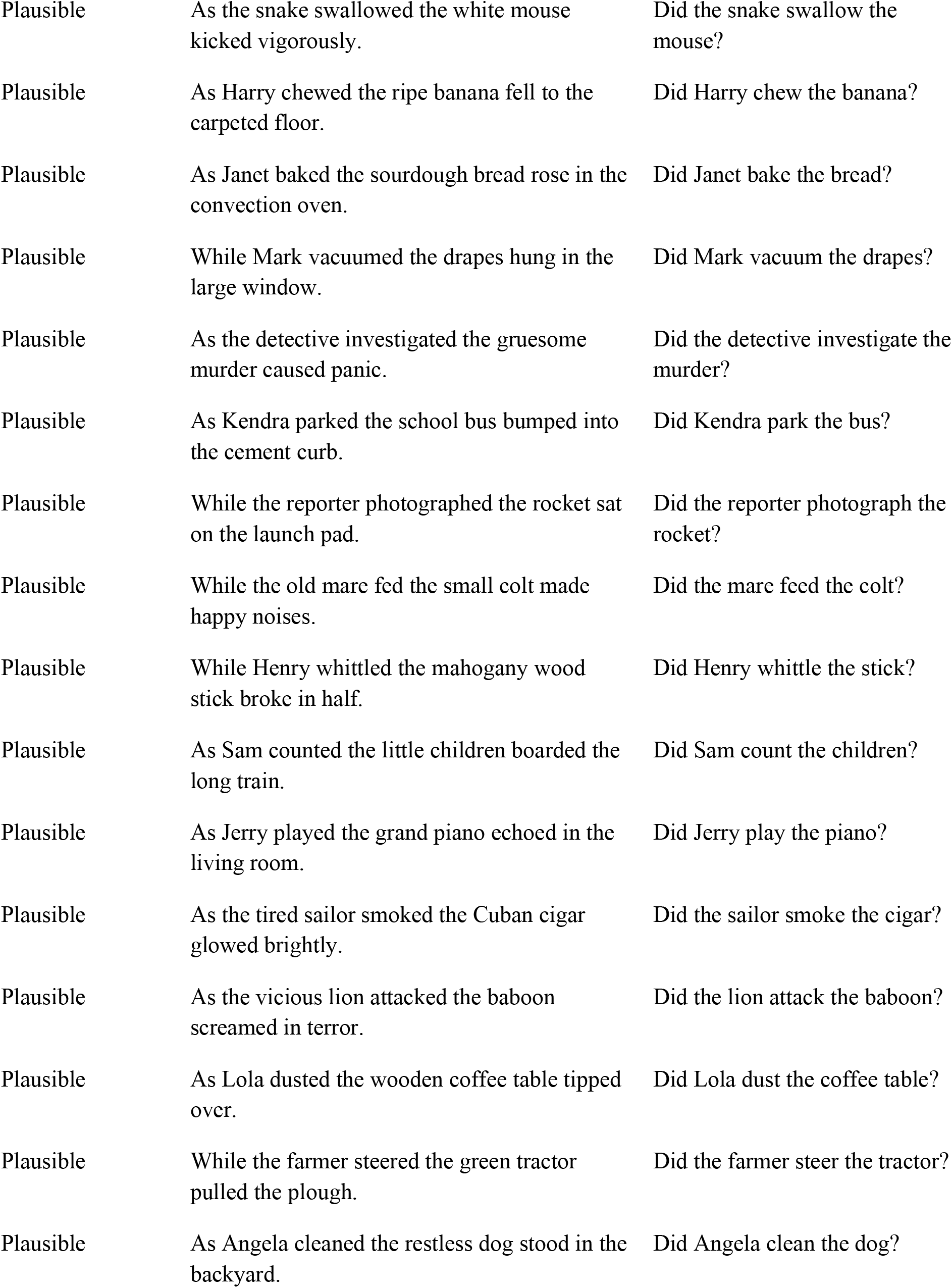
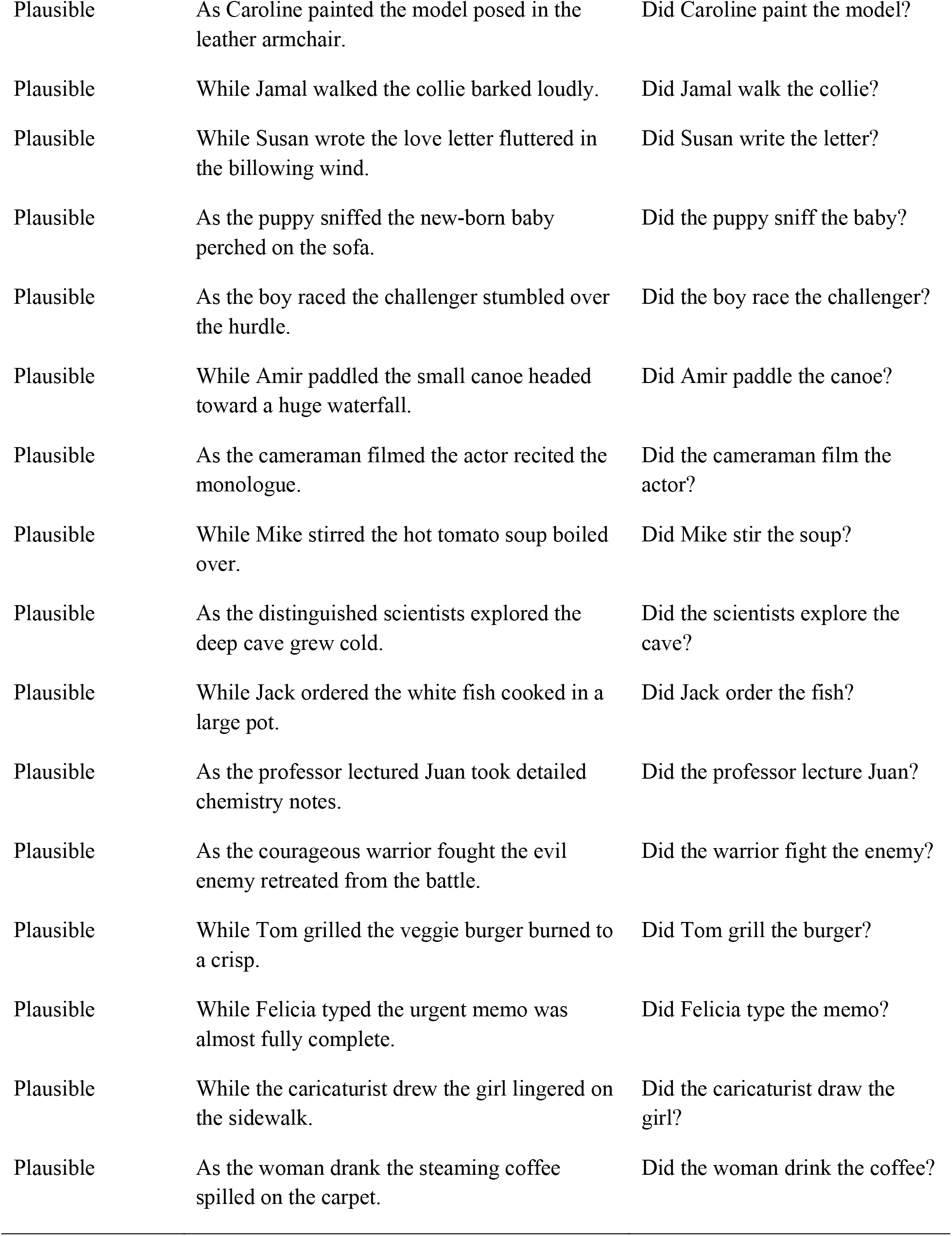
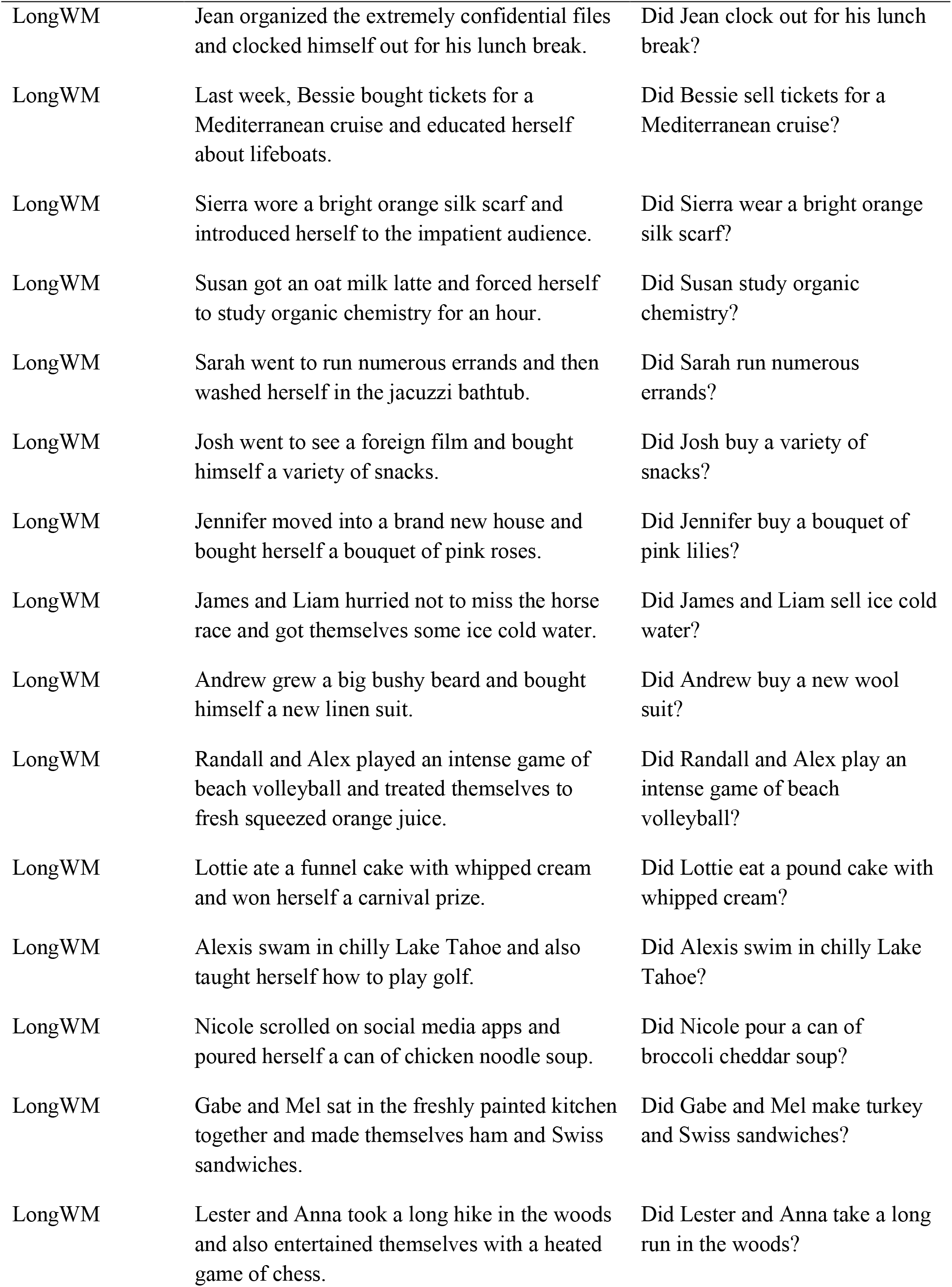
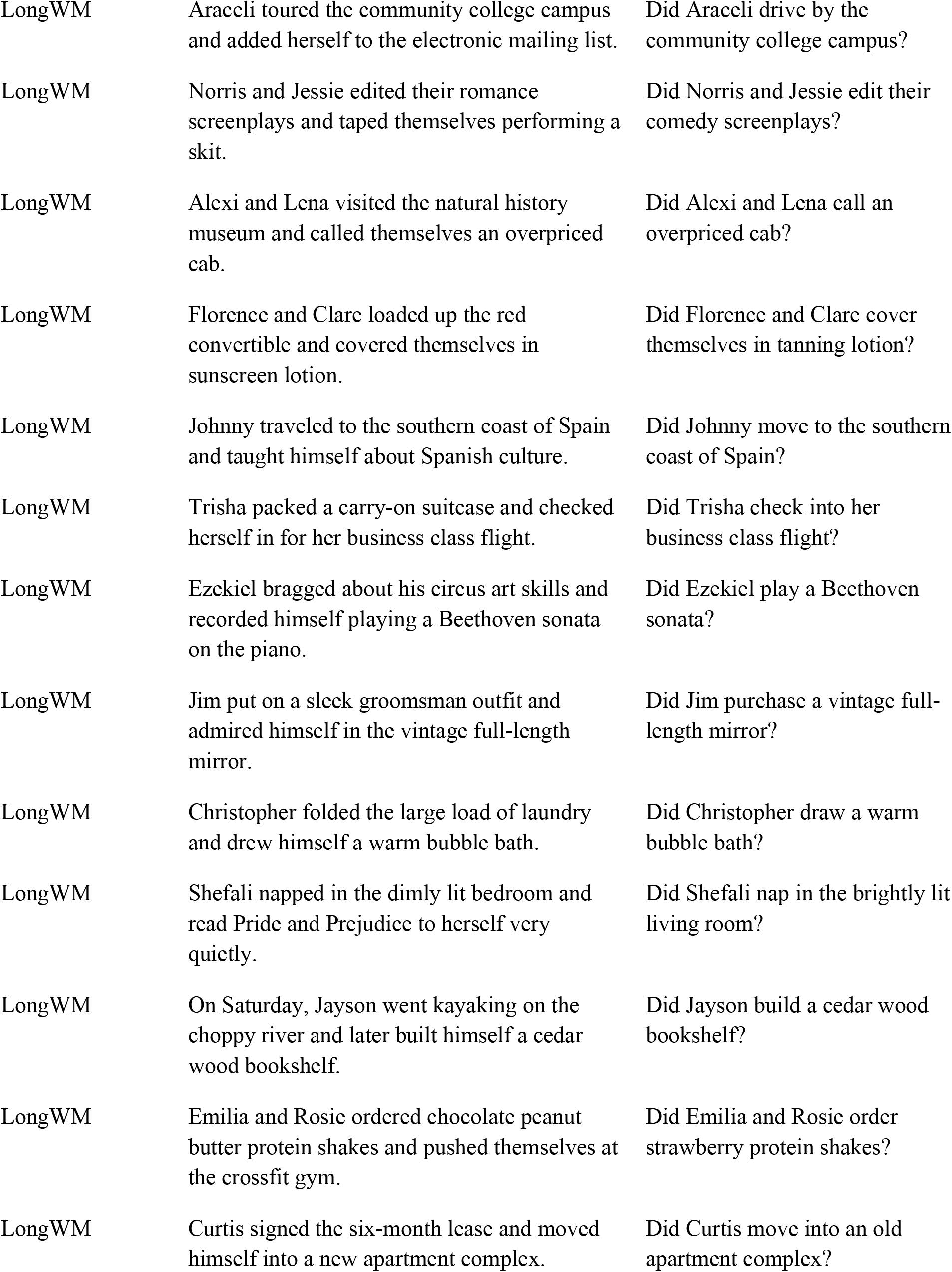
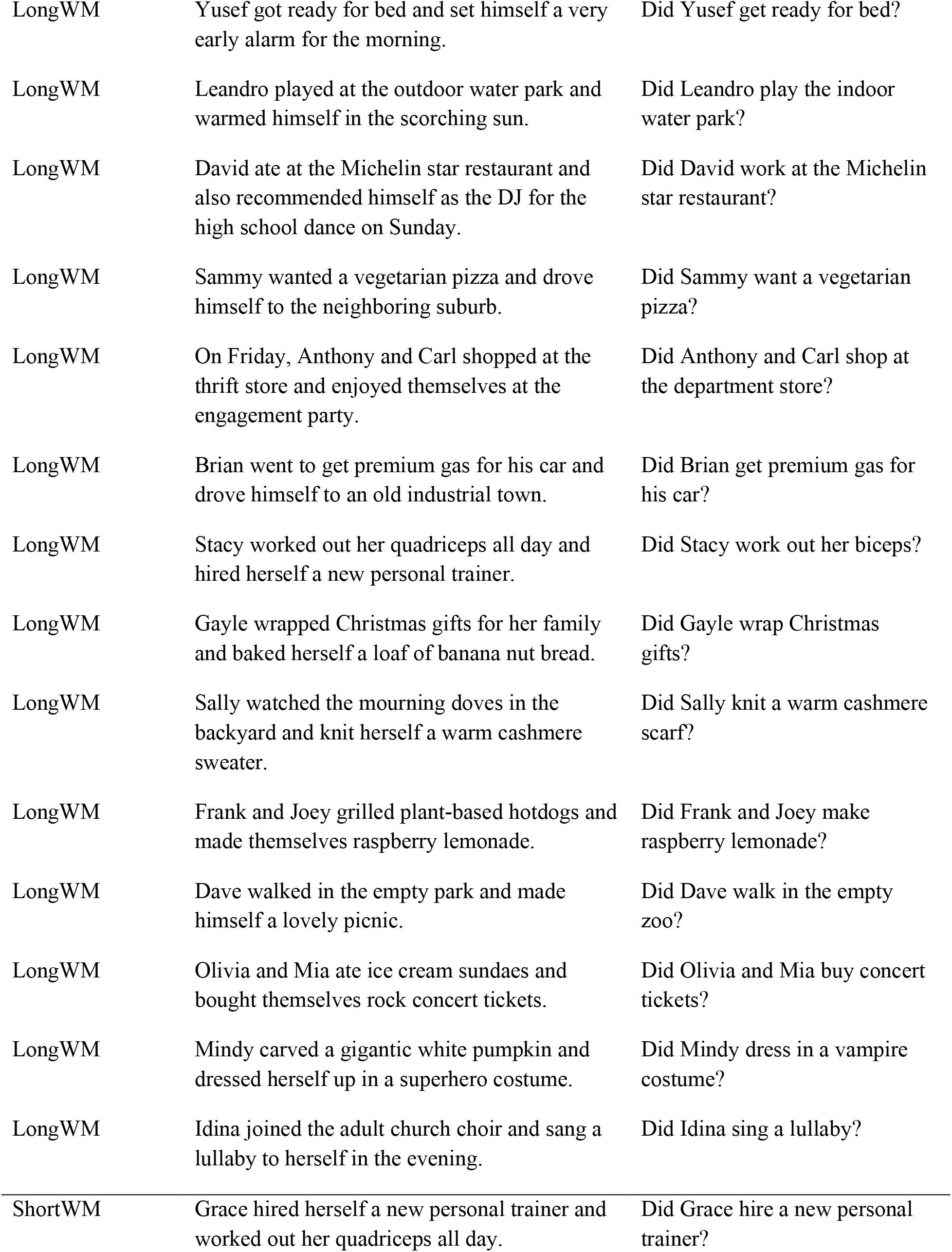
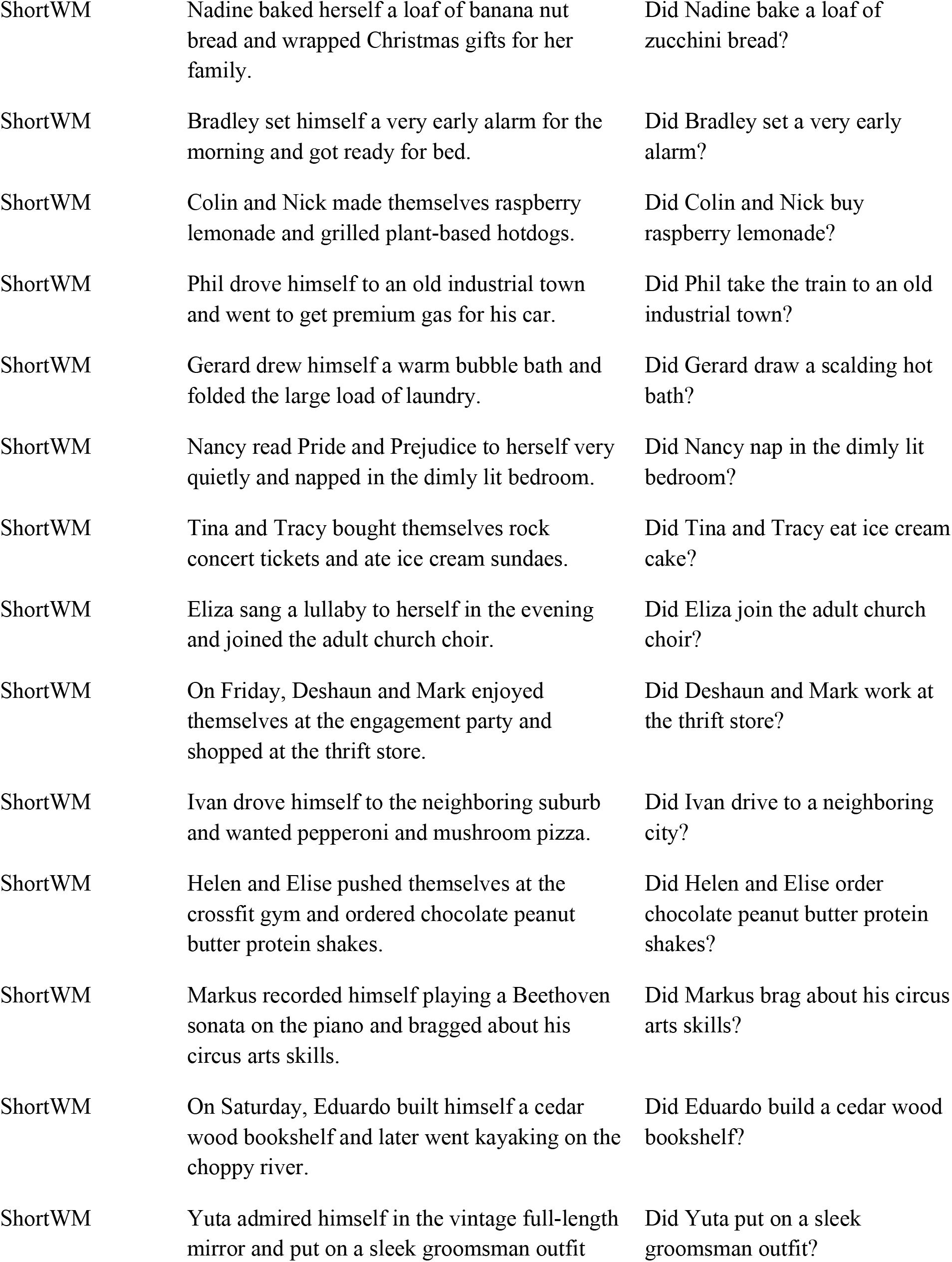
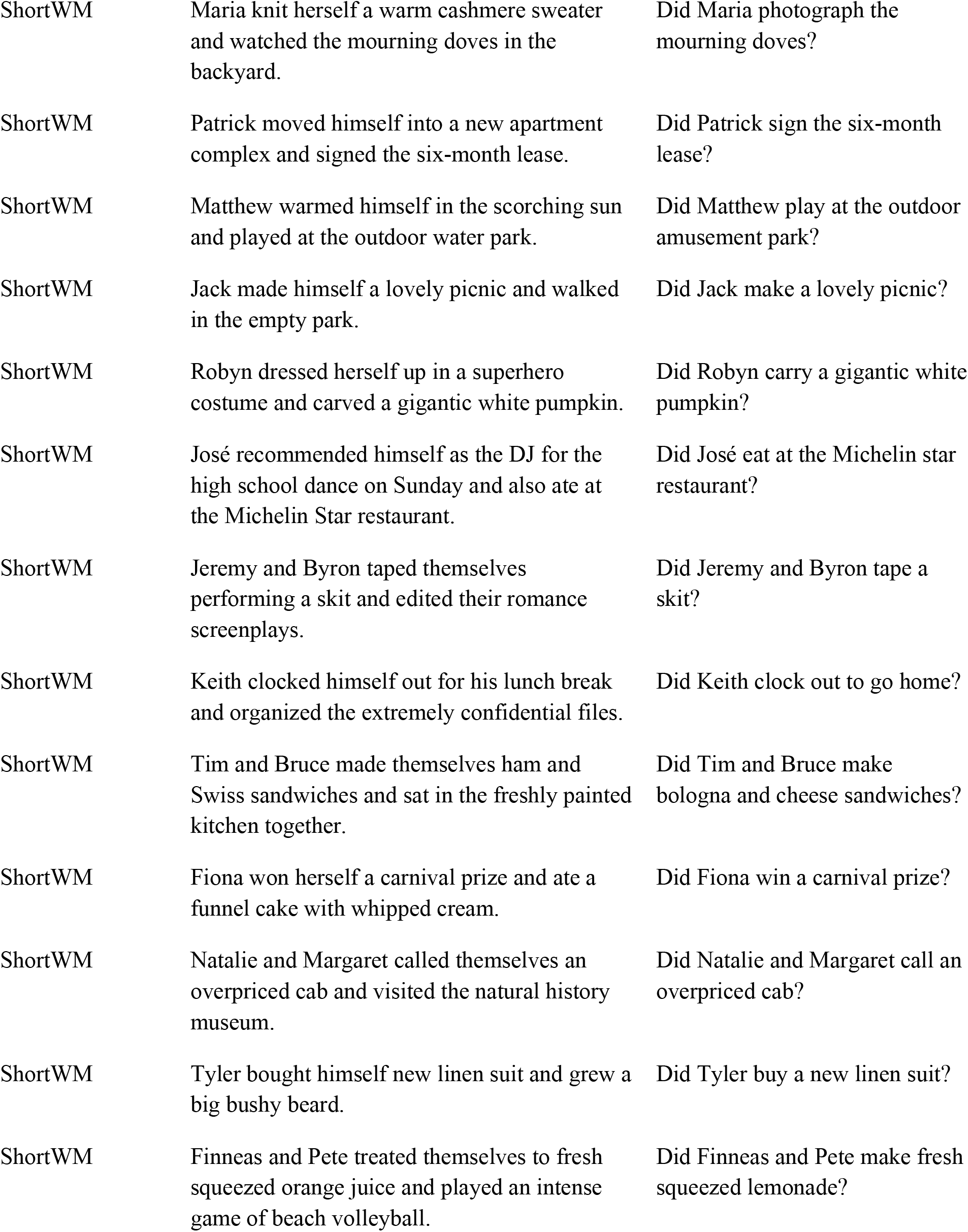
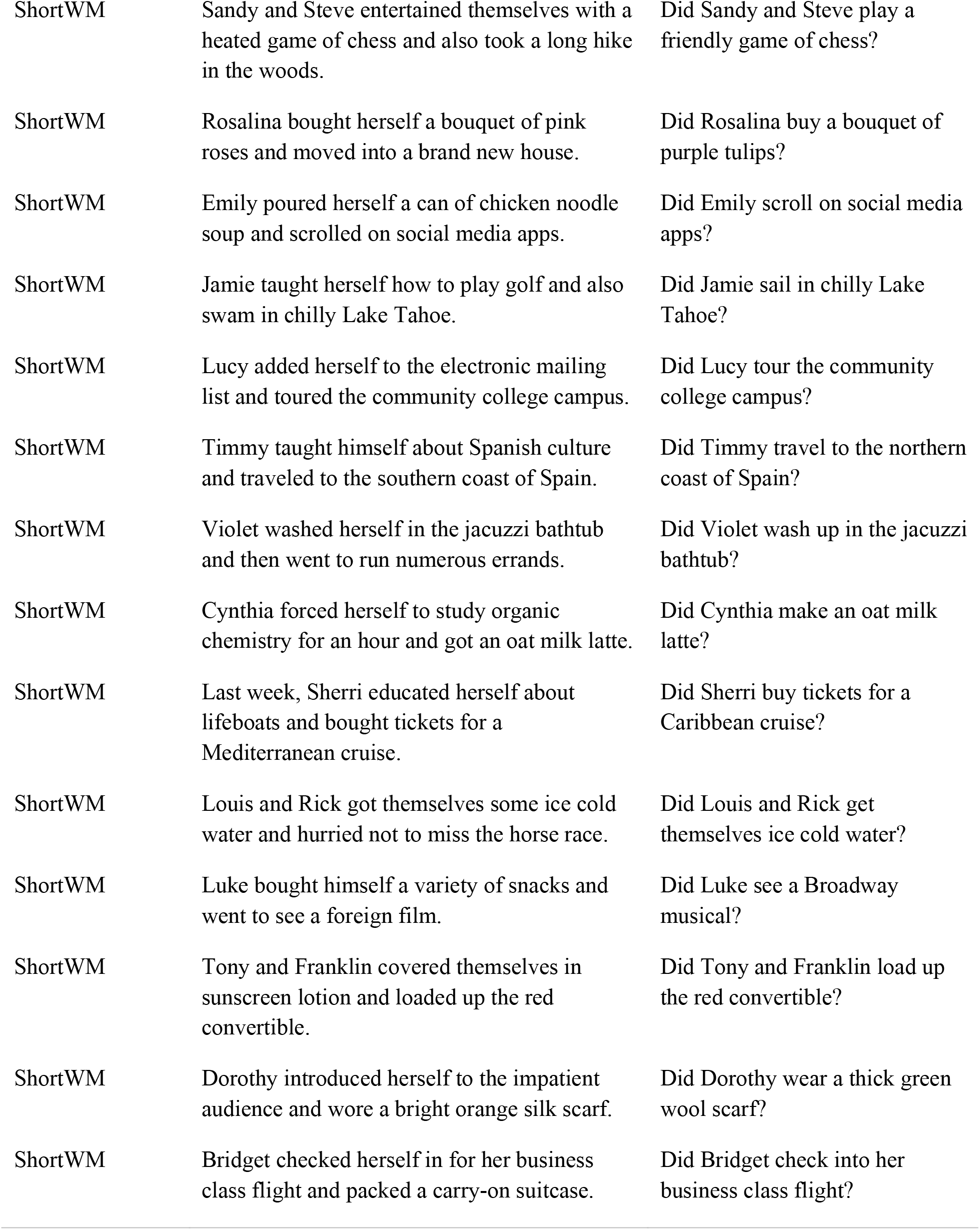

| Condition | Sentence | Question |
| --- | --- | --- |
| Implausible | While Harry steered the wooden canoe was moored at the dock. | Did Harry steer the canoe? |
| Implausible | While Henry stirred the sauce boiled on another burner. | Did Henry stir the sauce? |
| Implausible | As the girls raced the competitive opponents watched from the sideline. | Did the girls race the opponents? |
| Implausible | As the artist painted the self-portrait hung in the famous museum. | Did the artist paint the portrait? |
| Implausible | While the woman ordered the brown chicken escaped the farm. | Did the woman order the chicken? |
| Implausible | While Tom wrote the note rested in the trashcan with the waste. | Did Tom write the note? |
| Implausible | As the purring kitten sniffed the catnip was missing from the bag. | Did the kitten sniff the catnip? |
| Implausible | As the maid dusted the grand piano was left untouched and silent. | Did the maid dust the piano? |
| Implausible | As Rick explored the job opportunity escaped his notice entirely. | Did Rick explore the job opportunity? |
| Implausible | As the cameraman filmed the crew was nowhere to be found. | Did the cameraman film the crew? |
| Implausible | While the assistant typed the transcript came in the mail. | Did the assistant type the transcript? |
| Implausible | As Lola drank the white wine perched unopened in the wine cellar. | Did Lola drink the wine? |
| Implausible | While Janet walked the rambunctious pug played at home. | Did Janet walk the pug? |
| Implausible | As the soldiers fought the enemies never appeared for battle. | Did the soldiers fight the enemies? |
| Implausible | While Jerry drew the complicated blueprint was shipped overseas. | Did Jerry draw the blueprint? |
| Implausible | As Caroline cleaned the pan stayed in the oven with the vegetables. | Did Caroline clean the pan? |
| Implausible | While Angela grilled the chicken kebabs cooled in the fridge. | Did Angela grill the kebabs? |
| Implausible | While Jack paddled the kayak leaned in the shed untouched. | Did Jack paddle the kayak? |
| Implausible | As Sam smoked the hookah sat unused on the highest shelf. | Did Sam smoke the hookah? |
| Implausible | As the eagle ate the sparrow sheltered safe and sound. | Did the eagle eat the sparrow? |
| Implausible | As the teacher lectured the parents missed the meeting unknowingly. | Did the teacher lecture the parents? |
| Implausible | As the boys counted the small dogs were nowhere to be seen. | Did the boys count the dogs? |
| Implausible | While the cowboy rode the bull stood by the fence unfazed. | Did the cowboy ride the bull? |
| Implausible | While Kendra juggled the torches dangled high on the wall. | Did Kendra juggle the torches? |
| Implausible | While Felicia whittled the pencil lay unsharpened in the box. | Did Felicia whittle the pencil? |
| Implausible | While Juan drove the monster truck remained in the empty lot. | Did Juan drive the truck? |
| Implausible | While the student read the book burned in the fireplace. | Did the student read the book? |
| Implausible | As the police investigated the heist occurred undetected at the store. | Did the police investigate the heist? |
| Implausible | As the hooting owl swallowed the rat hid in the deep hole. | Did the owl swallow the rat? |
| Implausible | As the men wrestled the rivals arrived at the gymnasium. | Did the men wrestle the rivals? |
| Implausible | While the traveler sailed the dinghy stayed by the shore. | Did the traveler sail the dinghy? |
| Implausible | While the dancer performed the ballet was cancelled suddenly. | Did the dancer perform the ballet? |
| Implausible | While Bill hunted the lion paced in the zoo quickly. | Did Bill hunt the lion? |
| Implausible | As Mark ate the plump turkey roamed in its spacious cage. | Did Mark eat the turkey? |
| Implausible | While the intern photographed the workers hid from her. | Did the intern photograph the workers? |
| Implausible | While the brown cow fed the calf slept in the barn. | Did the brown cow feed the calf? |
| Implausible | While Susan studied the graphs resided unexamined on the board. | Did Susan study the graphs? |
| Implausible | As Klaus parked the sports car sped quickly away. | Did Klaus park the car? |
| Implausible | As Maria chewed the white bread laid uneaten in the food pantry. | Did Maria chew the bread? |
| Implausible | As Jamal baked the apple pie waited be made soon. | Did Jamal bake the pie? |
| Implausible | As Mike played the violin was placed in its case carefully. | Did Mike play the violin? |
| Implausible | While Amir vacuumed the dirty carpet spun in the dryer rapidly. | Did Amir vacuum the carpet? |
| Control | While Bill stirred the stew the soup boiled on the stove. | Did Bill stir the stew? |
| Control | While Jerry juggled the balls the props rested in the box. | Did Jerry juggle the balls? |
| Control | As the musician played the trumpet the oboe balanced on the stand. | Did the musician play the trumpet? |
| Control | While the zookeeper fed the ponies the stallion stomped its hoof. | Did the zookeeper feed the ponies? |
| Control | As Jamal cleaned the floor the dust blew away. | Did Jamal clean the floor? |
| Control | As the men parked the mopeds the tractor went in for repairs. | Did the men park the mopeds? |
| Control | As the teacher baked the cookies the cake was decorated. | Did the teacher bake the cookies? |
| Control | As the troops fought the invaders the civilians escaped to the shelter. | Did the troops fight the invaders? |
| Control | While Rick drove the limo the truck pulled over. | Did Rick drive the limo? |
| Control | As the woman wrestled the opponent the onlookers started yelling. | Did the woman wrestle the opponent? |
| Control | As Mark filmed the scene the actress took a break. | Did Mark film the scene? |
| Control | While the woman walked the poodle the pug slept in the house. | Did the woman walk the poodle? |
| Control | As the speaker lectured the audience the visitors returned home. | Did the speaker lecture the audience? |
| Control | While Janet wrote the comments the letter sat on the porch. | Did Janet write the comments? |
| Control | As Tom investigated the attack the suspect stayed in jail. | Did Tom investigate the attack? |
| Control | As the horse sniffed the hay the weeds swayed in the wind. | Did the horse sniff the hay? |
| Control | While Harry photographed the astronaut the probe landed on Mars. | Did Harry photograph the astronaut? |
| Control | As Sam painted the portrait the sculptor left for lunch. | Did Sam paint the portrait? |
| Control | As Mike drank the soda the tea brewed in the pot. | Did Mike drink the soda? |
| Control | While Angela sailed the schooner the jetski lingered at the dock. | Did Angela sail the schooner? |
| Control | While the women studied the diagram the report dangled on the bulletin board. | Did the women study the diagram? |
| Control | While Susan whittled the wood the twigs burned in the campfire. | Did Susan whittle the wood? |
| Control | While the students drew the landscape the vase held the flowers. | Did the students draw the landscape? |
| Control | While Amir steered the baler the cows chewed the grass. | Did Amir steer the baler? |
| Control | While the travelers rode the mules the camels drank water. | Did the travelers ride the mules? |
| Control | While the guests ordered the chicken the fish swam in the tank. | Did the guests order the chicken? |
| Control | As Henry counted the cars the girls formed a group. | Did Henry count the cars? |
| Control | As Klaus dusted the bookshelves the collectibles were clean. | Did Klaus dust the bookshelves? |
| Control | While Lola paddled the raft the canoe floated nearby. | Did Lola paddle the raft? |
| Control | As Kendra smoked the cigarette the cigar remained unlit. | Did Kendra smoke the cigarette? |
| Control | While Felicia grilled the steaks the ribs waited on the table. | Did Felicia grill the steaks? |
| Control | As Maria raced the competitor the team cheered in the stands. | Did Maria race the competitor? |
| Control | While the assistant typed the email the schedule remained the same. | Did the assistant type the email? |
| Control | As the workers ate the fruit the bread rose in the oven. | Did the workers eat the fruit? |
| Control | While Jack hunted the turkey the squirrel dashed into the bush. | Did Jack hunt the turkey? |
| Control | As Juan chewed the gum the meat thawed on the counter. | Did Juan chew the gum? |
| Control | As the hornets attacked the crowd the children ran to safety. | Did the hornets attack the crowd? |
| Control | While the boys vacuumed the stairs the rug lay on the couch. | Did the boys vacuum the stairs? |
| Control | While Caroline read the newspaper the documents lay in the drawer. | Did Caroline read the newspaper? |
| Control | While the Broadway stars performed the number the show impressed the patrons. | Did the Broadway stars perform the number? |
| Control | As the cat swallowed the lizard the birds flew away. | Did the cat swallow the lizard? |
| Control | As the hikers explored the forest the meadows were blooming with wildflowers. | Did the hikers explore the forest? |
| Plausible | While Rick drove the pick-up truck plunged into a ditch. | Did Rick drive the truck? |
| Plausible | As the athlete wrestled the tough opponent shouted insults. | Did the athlete wrestle the opponent? |
| Plausible | While the clown juggled the green apples dropped to the ground. | Did the clown juggle the apples? |
| Plausible | While Maria read the newspaper got wet from her hot coffee. | Did Maria read the newspaper? |
| Plausible | While the skipper sailed the cargo ship veered off course. | Did the skipper sail the cargo ship? |
| Plausible | While the rowdy cowboy rode the anxious horse neighed loudly. | Did the cowboy ride the horse? |
| Plausible | While the orchestra performed the symphony sounded beautiful. | Did the orchestra perform the symphony? |
| Plausible | While Klaus studied the chemistry textbook remained on the desk. | Did Klaus study the textbook? |
| Plausible | As Bill ate the ham lay on the dining room table. | Did Bill eat the ham? |
| Plausible | While the big man hunted the deer ran into the dark woods. | Did the big man hunt the deer? |
| Plausible | As the snake swallowed the white mouse kicked vigorously. | Did the snake swallow the mouse? |
| Plausible | As Harry chewed the ripe banana fell to the carpeted floor. | Did Harry chew the banana? |
| Plausible | As Janet baked the sourdough bread rose in the convection oven. | Did Janet bake the bread? |
| Plausible | While Mark vacuumed the drapes hung in the large window. | Did Mark vacuum the drapes? |
| Plausible | As the detective investigated the gruesome murder caused panic. | Did the detective investigate the murder? |
| Plausible | As Kendra parked the school bus bumped into the cement curb. | Did Kendra park the bus? |
| Plausible | While the reporter photographed the rocket sat on the launch pad. | Did the reporter photograph the rocket? |
| Plausible | While the old mare fed the small colt made happy noises. | Did the mare feed the colt? |
| Plausible | While Henry whittled the mahogany wood stick broke in half. | Did Henry whittle the stick? |
| Plausible | As Sam counted the little children boarded the long train. | Did Sam count the children? |
| Plausible | As Jerry played the grand piano echoed in the living room. | Did Jerry play the piano? |
| Plausible | As the tired sailor smoked the Cuban cigar glowed brightly. | Did the sailor smoke the cigar? |
| Plausible | As the vicious lion attacked the baboon screamed in terror. | Did the lion attack the baboon? |
| Plausible | As Lola dusted the wooden coffee table tipped over. | Did Lola dust the coffee table? |
| Plausible | While the farmer steered the green tractor pulled the plough. | Did the farmer steer the tractor? |
| Plausible | As Angela cleaned the restless dog stood in the backyard. | Did Angela clean the dog? |
| Plausible | As Caroline painted the model posed in the leather armchair. | Did Caroline paint the model? |
| Plausible | While Jamal walked the collie barked loudly. | Did Jamal walk the collie? |
| Plausible | While Susan wrote the love letter fluttered in the billowing wind. | Did Susan write the letter? |
| Plausible | As the puppy sniffed the new-born baby perched on the sofa. | Did the puppy sniff the baby? |
| Plausible | As the boy raced the challenger stumbled over the hurdle. | Did the boy race the challenger? |
| Plausible | While Amir paddled the small canoe headed toward a huge waterfall. | Did Amir paddle the canoe? |
| Plausible | As the cameraman filmed the actor recited the monologue. | Did the cameraman film the actor? |
| Plausible | While Mike stirred the hot tomato soup boiled over. | Did Mike stir the soup? |
| Plausible | As the distinguished scientists explored the deep cave grew cold. | Did the scientists explore the cave? |
| Plausible | While Jack ordered the white fish cooked in a large pot. | Did Jack order the fish? |
| Plausible | As the professor lectured Juan took detailed chemistry notes. | Did the professor lecture Juan? |
| Plausible | As the courageous warrior fought the evil enemy retreated from the battle. | Did the warrior fight the enemy? |
| Plausible | While Tom grilled the veggie burger burned to a crisp. | Did Tom grill the burger? |
| Plausible | While Felicia typed the urgent memo was almost fully complete. | Did Felicia type the memo? |
| Plausible | While the caricaturist drew the girl lingered on the sidewalk. | Did the caricaturist draw the girl? |
| Plausible | As the woman drank the steaming coffee spilled on the carpet. | Did the woman drink the coffee? |
| LongWM | Jean organized the extremely confidential files and clocked himself out for his lunch break. | Did Jean clock out for his lunch break? |
| LongWM | Last week, Bessie bought tickets for a Mediterranean cruise and educated herself about lifeboats. | Did Bessie sell tickets for a Mediterranean cruise? |
| LongWM | Sierra wore a bright orange silk scarf and introduced herself to the impatient audience. | Did Sierra wear a bright orange silk scarf? |
| LongWM | Susan got an oat milk latte and forced herself to study organic chemistry for an hour. | Did Susan study organic chemistry? |
| LongWM | Sarah went to run numerous errands and then washed herself in the jacuzzi bathtub. | Did Sarah run numerous errands? |
| LongWM | Josh went to see a foreign film and bought himself a variety of snacks. | Did Josh buy a variety of snacks? |
| LongWM | Jennifer moved into a brand new house and bought herself a bouquet of pink roses. | Did Jennifer buy a bouquet of pink lilies? |
| LongWM | James and Liam hurried not to miss the horse race and got themselves some ice cold water. | Did James and Liam sell ice cold water? |
| LongWM | Andrew grew a big bushy beard and bought himself a new linen suit. | Did Andrew buy a new wool suit? |
| LongWM | Randall and Alex played an intense game of beach volleyball and treated themselves to fresh squeezed orange juice. | Did Randall and Alex play an intense game of beach volleyball? |
| LongWM | Lottie ate a funnel cake with whipped cream and won herself a carnival prize. | Did Lottie eat a pound cake with whipped cream? |
| LongWM | Alexis swam in chilly Lake Tahoe and also taught herself how to play golf. | Did Alexis swim in chilly Lake Tahoe? |
| LongWM | Nicole scrolled on social media apps and poured herself a can of chicken noodle soup. | Did Nicole pour a can of broccoli cheddar soup? |
| LongWM | Gabe and Mel sat in the freshly painted kitchen together and made themselves ham and Swiss sandwiches. | Did Gabe and Mel make turkey and Swiss sandwiches? |
| LongWM | Lester and Anna took a long hike in the woods and also entertained themselves with a heated game of chess. | Did Lester and Anna take a long run in the woods? |
| LongWM | Araceli toured the community college campus and added herself to the electronic mailing list. | Did Araceli drive by the community college campus? |
| LongWM | Norris and Jessie edited their romance screenplays and taped themselves performing a skit. | Did Norris and Jessie edit their comedy screenplays? |
| LongWM | Alexi and Lena visited the natural history museum and called themselves an overpriced cab. | Did Alexi and Lena call an overpriced cab? |
| LongWM | Florence and Clare loaded up the red convertible and covered themselves in sunscreen lotion. | Did Florence and Clare cover themselves in tanning lotion? |
| LongWM | Johnny traveled to the southern coast of Spain and taught himself about Spanish culture. | Did Johnny move to the southern coast of Spain? |
| LongWM | Trisha packed a carry-on suitcase and checked herself in for her business class flight. | Did Trisha check into her business class flight? |
| LongWM | Ezekiel bragged about his circus art skills and recorded himself playing a Beethoven sonata on the piano. | Did Ezekiel play a Beethoven sonata? |
| LongWM | Jim put on a sleek groomsman outfit and admired himself in the vintage full-length mirror. | Did Jim purchase a vintage full-length mirror? |
| LongWM | Christopher folded the large load of laundry and drew himself a warm bubble bath. | Did Christopher draw a warm bubble bath? |
| LongWM | Shefali napped in the dimly lit bedroom and read Pride and Prejudice to herself very quietly. | Did Shefali nap in the brightly lit living room? |
| LongWM | On Saturday, Jayson went kayaking on the choppy river and later built himself a cedar wood bookshelf. | Did Jayson build a cedar wood bookshelf? |
| LongWM | Emilia and Rosie ordered chocolate peanut butter protein shakes and pushed themselves at the crossfit gym. | Did Emilia and Rosie order strawberry protein shakes? |
| LongWM | Curtis signed the six-month lease and moved himself into a new apartment complex. | Did Curtis move into an old apartment complex? |
| LongWM | Yusef got ready for bed and set himself a very early alarm for the morning. | Did Yusef get ready for bed? |
| LongWM | Leandro played at the outdoor water park and warmed himself in the scorching sun. | Did Leandro play the indoor water park? |
| LongWM | David ate at the Michelin star restaurant and also recommended himself as the DJ for the high school dance on Sunday. | Did David work at the Michelin star restaurant? |
| LongWM | Sammy wanted a vegetarian pizza and drove himself to the neighboring suburb. | Did Sammy want a vegetarian pizza? |
| LongWM | On Friday, Anthony and Carl shopped at the thrift store and enjoyed themselves at the engagement party. | Did Anthony and Carl shop at the department store? |
| LongWM | Brian went to get premium gas for his car and drove himself to an old industrial town. | Did Brian get premium gas for his car? |
| LongWM | Stacy worked out her quadriceps all day and hired herself a new personal trainer. | Did Stacy work out her biceps? |
| LongWM | Gayle wrapped Christmas gifts for her family and baked herself a loaf of banana nut bread. | Did Gayle wrap Christmas gifts? |
| LongWM | Sally watched the mourning doves in the backyard and knit herself a warm cashmere sweater. | Did Sally knit a warm cashmere scarf? |
| LongWM | Frank and Joey grilled plant-based hotdogs and made themselves raspberry lemonade. | Did Frank and Joey make raspberry lemonade? |
| LongWM | Dave walked in the empty park and made himself a lovely picnic. | Did Dave walk in the empty zoo? |
| LongWM | Olivia and Mia ate ice cream sundaes and bought themselves rock concert tickets. | Did Olivia and Mia buy concert tickets? |
| LongWM | Mindy carved a gigantic white pumpkin and dressed herself up in a superhero costume. | Did Mindy dress in a vampire costume? |
| LongWM | Idina joined the adult church choir and sang a lullaby to herself in the evening. | Did Idina sing a lullaby? |
| ShortWM | Grace hired herself a new personal trainer and worked out her quadriceps all day. | Did Grace hire a new personal trainer? |
| ShortWM | Nadine baked herself a loaf of banana nut bread and wrapped Christmas gifts for her family. | Did Nadine bake a loaf of zucchini bread? |
| ShortWM | Bradley set himself a very early alarm for the morning and got ready for bed. | Did Bradley set a very early alarm? |
| ShortWM | Colin and Nick made themselves raspberry lemonade and grilled plant-based hotdogs. | Did Colin and Nick buy raspberry lemonade? |
| ShortWM | Phil drove himself to an old industrial town and went to get premium gas for his car. | Did Phil take the train to an old industrial town? |
| ShortWM | Gerard drew himself a warm bubble bath and folded the large load of laundry. | Did Gerard draw a scalding hot bath? |
| ShortWM | Nancy read Pride and Prejudice to herself very quietly and napped in the dimly lit bedroom. | Did Nancy nap in the dimly lit bedroom? |
| ShortWM | Tina and Tracy bought themselves rock concert tickets and ate ice cream sundaes. | Did Tina and Tracy eat ice cream cake? |
| ShortWM | Eliza sang a lullaby to herself in the evening and joined the adult church choir. | Did Eliza join the adult church choir? |
| ShortWM | On Friday, Deshaun and Mark enjoyed themselves at the engagement party and shopped at the thrift store. | Did Deshaun and Mark work at the thrift store? |
| ShortWM | Ivan drove himself to the neighboring suburb and wanted pepperoni and mushroom pizza. | Did Ivan drive to a neighboring city? |
| ShortWM | Helen and Elise pushed themselves at the crossfit gym and ordered chocolate peanut butter protein shakes. | Did Helen and Elise order chocolate peanut butter protein shakes? |
| ShortWM | Markus recorded himself playing a Beethoven sonata on the piano and bragged about his circus arts skills. | Did Markus brag about his circus arts skills? |
| ShortWM | On Saturday, Eduardo built himself a cedar wood bookshelf and later went kayaking on the choppy river. | Did Eduardo build a cedar wood bookshelf? |
| ShortWM | Yuta admired himself in the vintage full-length mirror and put on a sleek groomsman outfit | Did Yuta put on a sleek groomsman outfit? |
| ShortWM | Maria knit herself a warm cashmere sweater and watched the mourning doves in the backyard. | Did Maria photograph the mourning doves? |
| ShortWM | Patrick moved himself into a new apartment complex and signed the six-month lease. | Did Patrick sign the six-month lease? |
| ShortWM | Matthew warmed himself in the scorching sun and played at the outdoor water park. | Did Matthew play at the outdoor amusement park? |
| ShortWM | Jack made himself a lovely picnic and walked in the empty park. | Did Jack make a lovely picnic? |
| ShortWM | Robyn dressed herself up in a superhero costume and carved a gigantic white pumpkin. | Did Robyn carry a gigantic white pumpkin? |
| ShortWM | José recommended himself as the DJ for the high school dance on Sunday and also ate at the Michelin Star restaurant. | Did José eat at the Michelin star restaurant? |
| ShortWM | Jeremy and Byron taped themselves performing a skit and edited their romance screenplays. | Did Jeremy and Byron tape a skit? |
| ShortWM | Keith clocked himself out for his lunch break and organized the extremely confidential files. | Did Keith clock out to go home? |
| ShortWM | Tim and Bruce made themselves ham and Swiss sandwiches and sat in the freshly painted kitchen together. | Did Tim and Bruce make bologna and cheese sandwiches? |
| ShortWM | Fiona won herself a carnival prize and ate a funnel cake with whipped cream. | Did Fiona win a carnival prize? |
| ShortWM | Natalie and Margaret called themselves an overpriced cab and visited the natural history museum. | Did Natalie and Margaret call an overpriced cab? |
| ShortWM | Tyler bought himself new linen suit and grew a big bushy beard. | Did Tyler buy a new linen suit? |
| ShortWM | Finneas and Pete treated themselves to fresh squeezed orange juice and played an intense game of beach volleyball. | Did Finneas and Pete make fresh squeezed lemonade? |
| ShortWM | Sandy and Steve entertained themselves with a heated game of chess and also took a long hike in the woods. | Did Sandy and Steve play a friendly game of chess? |
| ShortWM | Rosalina bought herself a bouquet of pink roses and moved into a brand new house. | Did Rosalina buy a bouquet of purple tulips? |
| ShortWM | Emily poured herself a can of chicken noodle soup and scrolled on social media apps. | Did Emily scroll on social media apps? |
| ShortWM | Jamie taught herself how to play golf and also swam in chilly Lake Tahoe. | Did Jamie sail in chilly Lake Tahoe? |
| ShortWM | Lucy added herself to the electronic mailing list and toured the community college campus. | Did Lucy tour the community college campus? |
| ShortWM | Timmy taught himself about Spanish culture and traveled to the southern coast of Spain. | Did Timmy travel to the northern coast of Spain? |
| ShortWM | Violet washed herself in the jacuzzi bathtub and then went to run numerous errands. | Did Violet wash up in the jacuzzi bathtub? |
| ShortWM | Cynthia forced herself to study organic chemistry for an hour and got an oat milk latte. | Did Cynthia make an oat milk latte? |
| ShortWM | Last week, Sherri educated herself about lifeboats and bought tickets for a Mediterranean cruise. | Did Sherri buy tickets for a Caribbean cruise? |
| ShortWM | Louis and Rick got themselves some ice cold water and hurried not to miss the horse race. | Did Louis and Rick get themselves ice cold water? |
| ShortWM | Luke bought himself a variety of snacks and went to see a foreign film. | Did Luke see a Broadway musical? |
| ShortWM | Tony and Franklin covered themselves in sunscreen lotion and loaded up the red convertible. | Did Tony and Franklin load up the red convertible? |
| ShortWM | Dorothy introduced herself to the impatient audience and wore a bright orange silk scarf. | Did Dorothy wear a thick green wool scarf? |
| ShortWM | Bridget checked herself in for her business class flight and packed a carry-on suitcase. | Did Bridget check into her business class flight? |

### Attention Stimuli

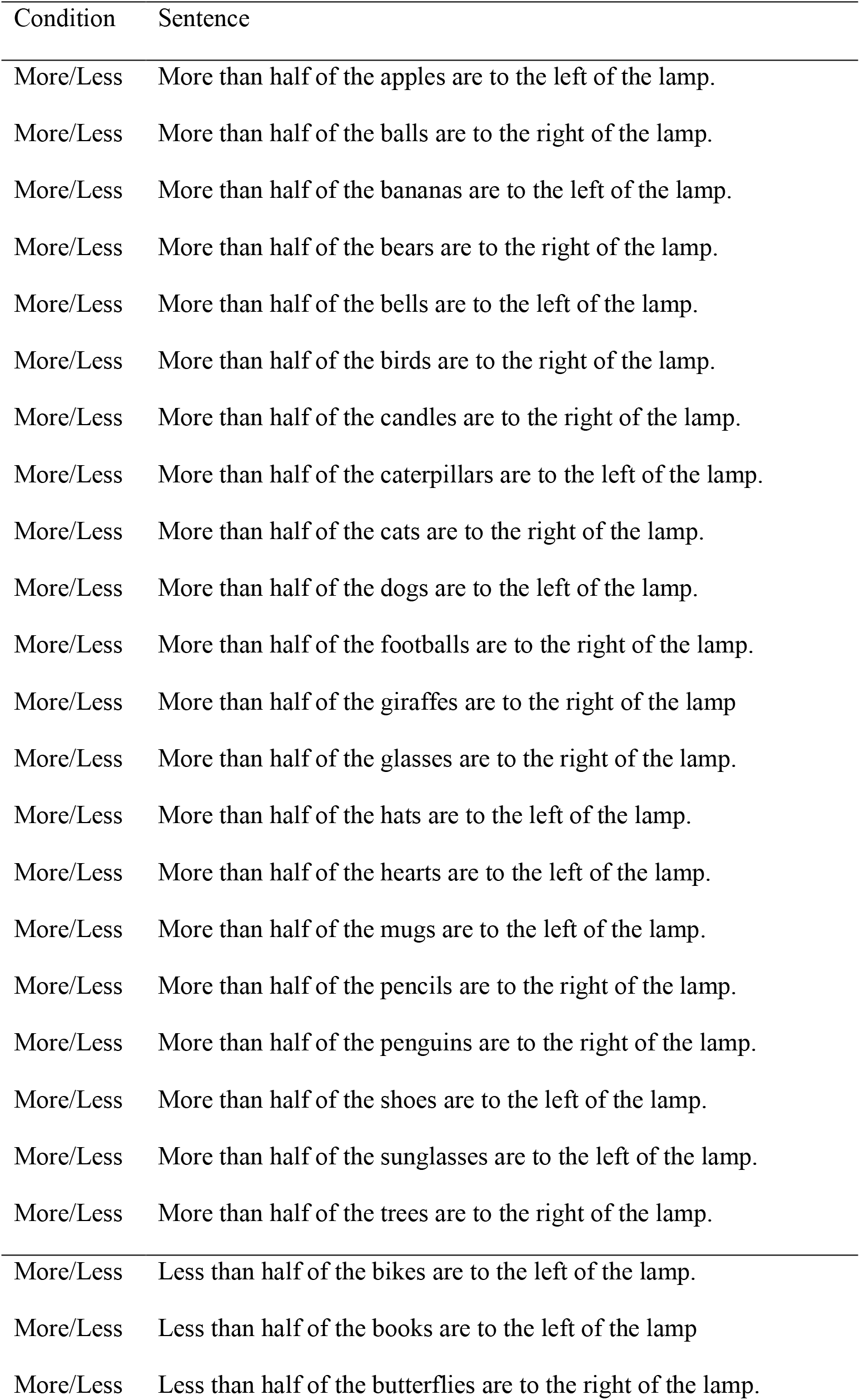
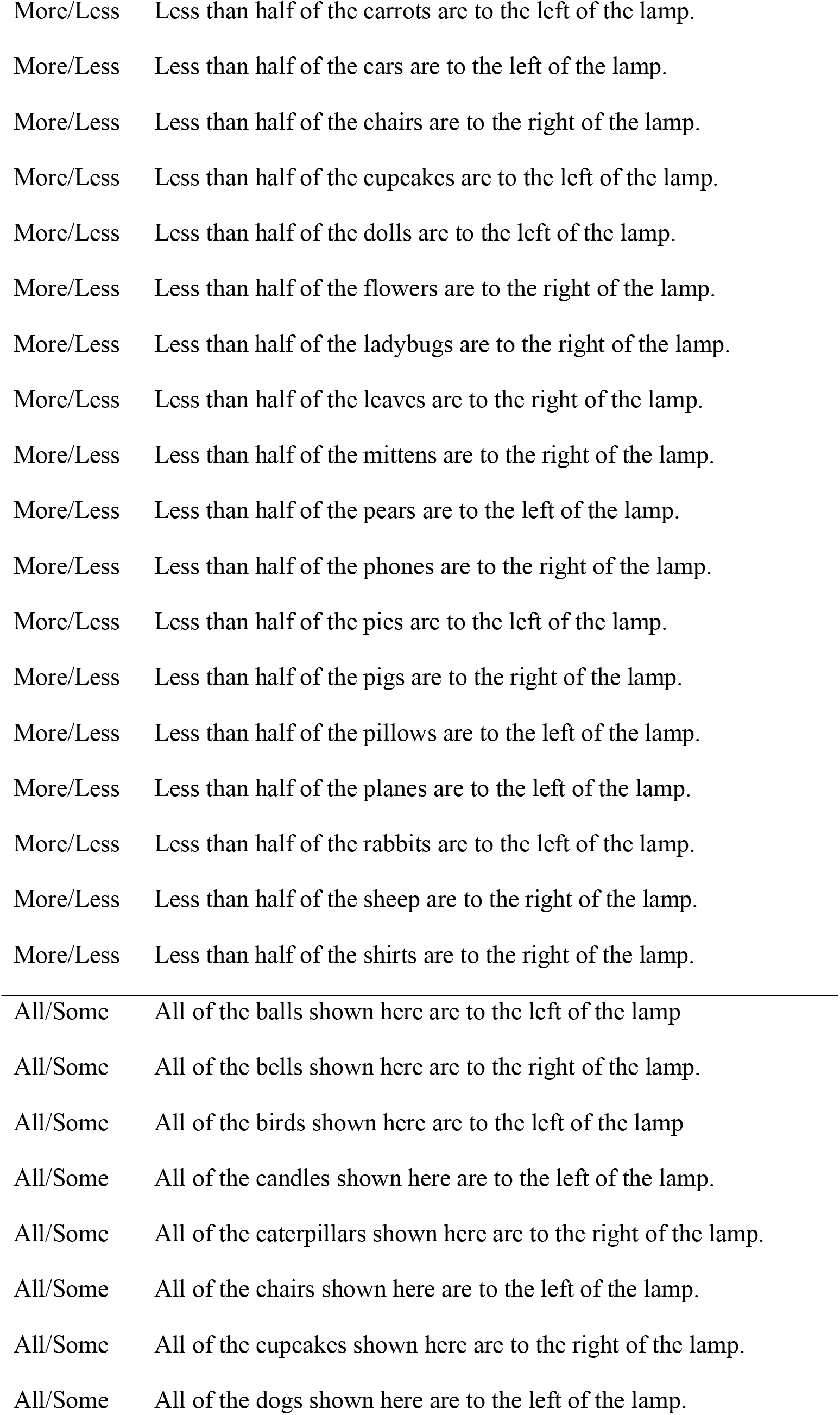
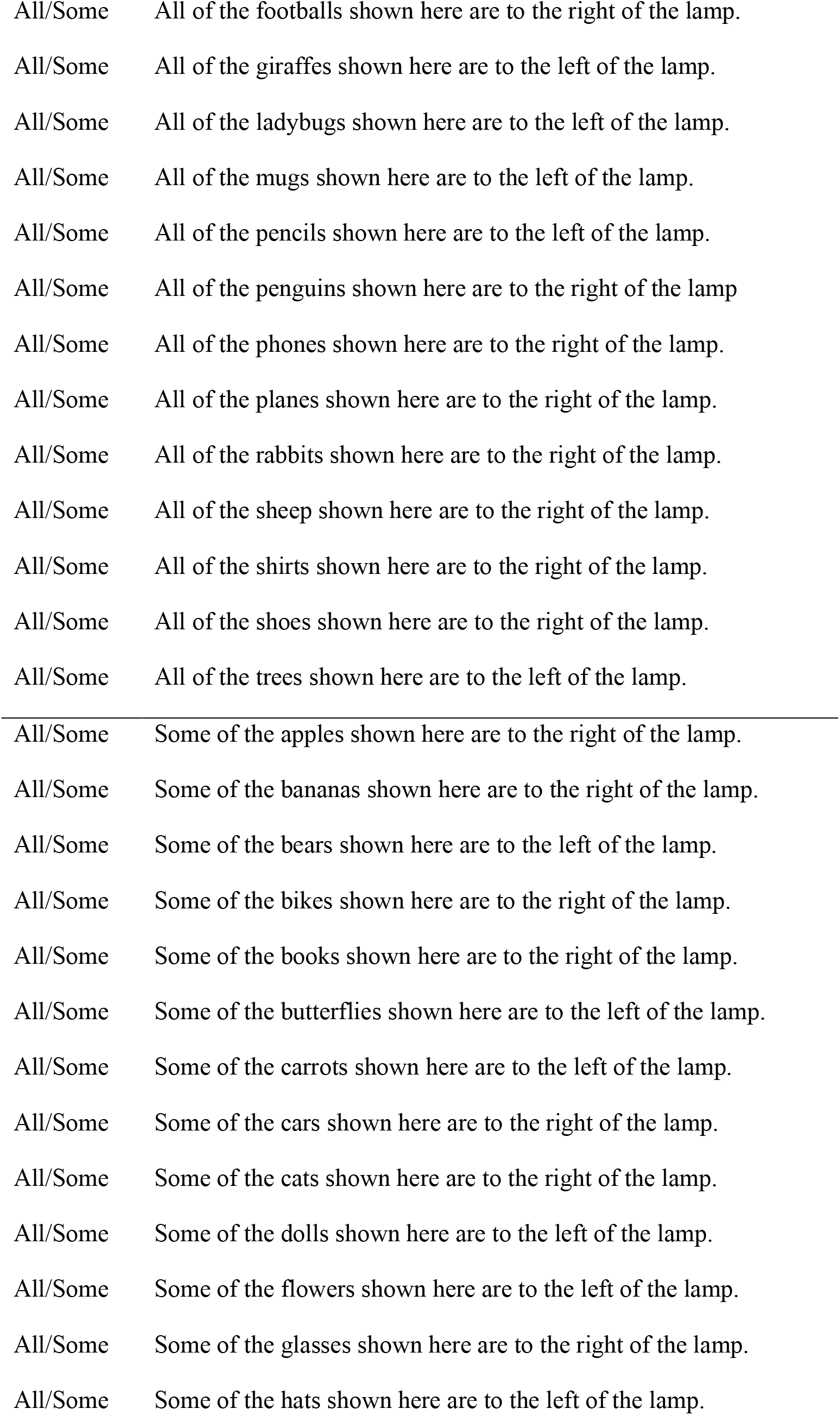
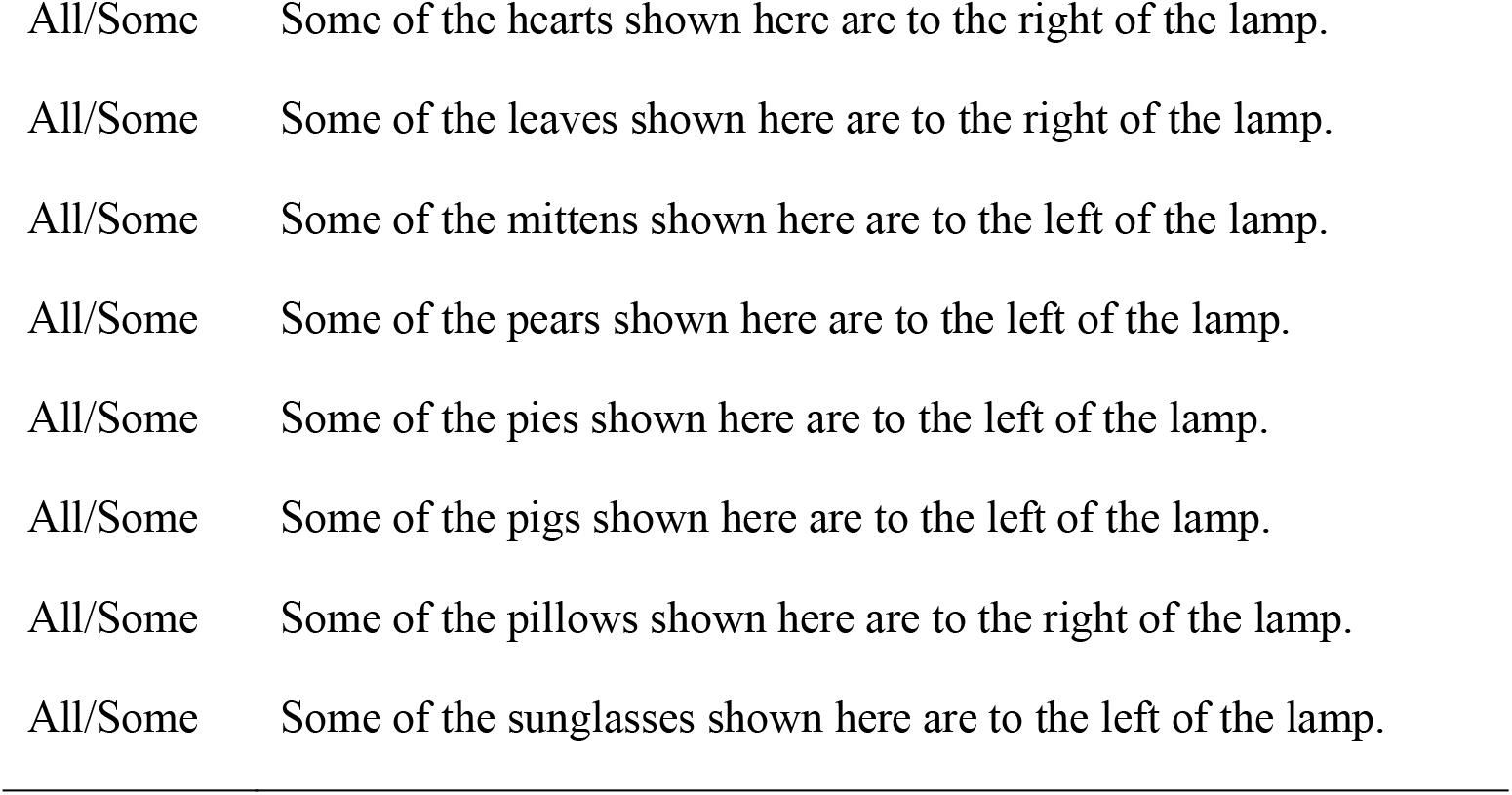

| Condition | Sentence |
| --- | --- |
| More/Less | More than half of the apples are to the left of the lamp. |
| More/Less | More than half of the balls are to the right of the lamp. |
| More/Less | More than half of the bananas are to the left of the lamp. |
| More/Less | More than half of the bears are to the right of the lamp. |
| More/Less | More than half of the bells are to the left of the lamp. |
| More/Less | More than half of the birds are to the right of the lamp. |
| More/Less | More than half of the candles are to the right of the lamp. |
| More/Less | More than half of the caterpillars are to the left of the lamp. |
| More/Less | More than half of the cats are to the right of the lamp. |
| More/Less | More than half of the dogs are to the left of the lamp. |
| More/Less | More than half of the footballs are to the right of the lamp. |
| More/Less | More than half of the giraffes are to the right of the lamp. |
| More/Less | More than half of the glasses are to the right of the lamp. |
| More/Less | More than half of the hats are to the left of the lamp. |
| More/Less | More than half of the hearts are to the left of the lamp. |
| More/Less | More than half of the mugs are to the left of the lamp. |
| More/Less | More than half of the pencils are to the right of the lamp. |
| More/Less | More than half of the penguins are to the right of the lamp. |
| More/Less | More than half of the shoes are to the left of the lamp. |
| More/Less | More than half of the sunglasses are to the left of the lamp. |
| More/Less | More than half of the trees are to the right of the lamp. |
| More/Less | Less than half of the bikes are to the left of the lamp. |
| More/Less | Less than half of the books are to the left of the lamp. |
| More/Less | Less than half of the butterflies are to the right of the lamp. |
| More/Less | Less than half of the carrots are to the left of the lamp. |
| More/Less | Less than half of the cars are to the left of the lamp. |
| More/Less | Less than half of the chairs are to the right of the lamp. |
| More/Less | Less than half of the cupcakes are to the left of the lamp. |
| More/Less | Less than half of the dolls are to the left of the lamp. |
| More/Less | Less than half of the flowers are to the right of the lamp. |
| More/Less | Less than half of the ladybugs are to the right of the lamp. |
| More/Less | Less than half of the leaves are to the right of the lamp. |
| More/Less | Less than half of the mittens are to the right of the lamp. |
| More/Less | Less than half of the pears are to the left of the lamp. |
| More/Less | Less than half of the phones are to the right of the lamp. |
| More/Less | Less than half of the pies are to the left of the lamp. |
| More/Less | Less than half of the pigs are to the right of the lamp. |
| More/Less | Less than half of the pillows are to the left of the lamp. |
| More/Less | Less than half of the planes are to the left of the lamp. |
| More/Less | Less than half of the rabbits are to the left of the lamp. |
| More/Less | Less than half of the sheep are to the right of the lamp. |
| More/Less | Less than half of the shirts are to the right of the lamp. |
| All/Some | All of the balls shown here are to the left of the lamp |
| All/Some | All of the bells shown here are to the right of the lamp. |
| All/Some | All of the birds shown here are to the left of the lamp |
| All/Some | All of the candles shown here are to the left of the lamp. |
| All/Some | All of the caterpillars shown here are to the right of the lamp. |
| All/Some | All of the chairs shown here are to the left of the lamp. |
| All/Some | All of the cupcakes shown here are to the right of the lamp. |
| All/Some | All of the dogs shown here are to the left of the lamp. |
| All/Some | All of the footballs shown here are to the right of the lamp. |
| All/Some | All of the giraffes shown here are to the left of the lamp. |
| All/Some | All of the ladybugs shown here are to the left of the lamp. |
| All/Some | All of the mugs shown here are to the left of the lamp. |
| All/Some | All of the pencils shown here are to the left of the lamp. |
| All/Some | All of the penguins shown here are to the right of the lamp |
| All/Some | All of the phones shown here are to the right of the lamp. |
| All/Some | All of the planes shown here are to the right of the lamp. |
| All/Some | All of the rabbits shown here are to the right of the lamp. |
| All/Some | All of the sheep shown here are to the right of the lamp. |
| All/Some | All of the shirts shown here are to the right of the lamp. |
| All/Some | All of the shoes shown here are to the right of the lamp. |
| All/Some | All of the trees shown here are to the left of the lamp. |
| All/Some | Some of the apples shown here are to the right of the lamp. |
| All/Some | Some of the bananas shown here are to the right of the lamp. |
| All/Some | Some of the bears shown here are to the left of the lamp. |
| All/Some | Some of the bikes shown here are to the right of the lamp. |
| All/Some | Some of the books shown here are to the right of the lamp. |
| All/Some | Some of the butterflies shown here are to the left of the lamp. |
| All/Some | Some of the carrots shown here are to the left of the lamp. |
| All/Some | Some of the cars shown here are to the right of the lamp. |
| All/Some | Some of the cats shown here are to the right of the lamp. |
| All/Some | Some of the dolls shown here are to the left of the lamp. |
| All/Some | Some of the flowers shown here are to the left of the lamp. |
| All/Some | Some of the glasses shown here are to the right of the lamp. |
| All/Some | Some of the hats shown here are to the left of the lamp. |
| All/Some | Some of the hearts shown here are to the right of the lamp. |
| All/Some | Some of the leaves shown here are to the right of the lamp. |
| All/Some | Some of the mittens shown here are to the left of the lamp. |
| All/Some | Some of the pears shown here are to the left of the lamp. |
| All/Some | Some of the pies shown here are to the left of the lamp. |
| All/Some | Some of the pigs shown here are to the left of the lamp. |
| All/Some | Some of the pillows shown here are to the right of the lamp. |
| All/Some | Some of the sunglasses shown here are to the left of the lamp. |

## Acknowledgments

This work was supported by a National Institutes of Health grant (R01DC017138) awarded to Malathi Thothathiri. A.G.L. was supported by Gravitation Grant 024.001.006 of the Language in Interaction Consortium from the Netherlands Organization for Scientific Research. A huge thanks to Luca Cavan and Phoebe Jeng for assistance with stimulus design, norming, and data collection.

## Footnotes

1 Due to a technical error, demographic information was lost for 7 adults. The reported numbers are for 30 out of 37 participants.

2 Sentences and questions were separated by variable jitters and modeled separately (see more below). We draw inferences based on neural activation for the sentences only. Therefore, differences in correct responses to the questions do not impact interpretation of the relevant fMRI results.

3 There is debate about whether null-hypothesis significance tests of confound variables are appropriate to conduct and interpret (Sassenhagen & Alday, 2016). For informational purposes, pairwise 2-tailed t-tests comparing the lengths of Implausible, Plausible and Control sentences did not reveal any significant differences (p’s>.05). Frequency comparisons between Implausible and Plausible sentences also did not reveal any significant differences (p’s>.05).

4 We used the peaks for Implausible versus Baseline rather than Implausible versus Control here because the former is orthogonal with the planned contrasts (Plausible vs Control, Long WM vs Control, and Short WM vs Control) and the latter is not. Using peaks identified using the latter would bias the ROIs towards voxels with low Control activation and thereby bias in favor of finding a difference of all conditions against Control.

