## Supplementary Info for "Fractionating difficulty during sentence comprehension using functional neuroimaging"

Supplementary Information

*SI #1: Suprathreshold clusters from the contrast of Implausible > Control*

| Cluster Index | # Voxels | Max Z value | Max Z coordinate (MNI) | | |
|---|---|---|---|---|---|
| 4 | 1498 | 5.37 | -52 | -42 | 6 |
| 3 | 738 | 5.14 | -50 | 28 | 0 |
| 2 | 155 | 4.57 | 52 | 22 | 22 |
| 1 | 105 | 4.14 | -4 | 46 | 42 |

*SI #2: Suprathreshold clusters from the contrast of Long WM > Control*

| Cluster Index | # Voxels | Max Z value | Max Z coordinate (MNI) | | |
|---|---|---|---|---|---|
| 8 | 7483 | 6.66 | 26 | -86 | 10 |
| 7 | 524 | 4.79 | -20 | -8 | -12 |
| 6 | 234 | 4.6 | -56 | -6 | -18 |
| 5 | 149 | 5.03 | -2 | -32 | 38 |
| 4 | 135 | 4.41 | -8 | 56 | 24 |
| 3 | 123 | 4.33 | -10 | 48 | 40 |
| 2 | 96 | 4.38 | -58 | -6 | -2 |
| 1 | 94 | 3.86 | -10 | 42 | 2 |
